# Evolution of plasmid mobility: origin and fate of non-conjugative plasmids

**DOI:** 10.1101/2021.12.10.472114

**Authors:** Charles Coluzzi, Maria del Pilar Garcillán-Barcia, Fernando de la Cruz, Eduardo P.C. Rocha

## Abstract

Conjugation drives horizontal gene transfer of many adaptive traits across prokaryotes. Yet, only a fourth of the plasmids encode the functions necessary to conjugate autonomously, others being non-mobile or mobilizable by other elements. How these different plasmids evolve is poorly understood. Here, we studied plasmid evolution in terms of their gene repertoires and relaxases. We observed that gene content in plasmid varies rapidly in relation to the rate of evolution of relaxases, such that plasmids with 95% identical relaxases have on average fewer than 50% of homologs. The identification of 249 recent transitions in terms of mobility types revealed that they are associated with even greater changes in gene repertoires, possibly mediated by transposable elements that are more abundant in such plasmids. These changes include pseudogenization of the conjugation locus, exchange of replication initiators, and extensive gene loss. In some instances, the transition between mobility types also leads to the genesis of novel plasmid taxonomic units. Most of these transitions are short-lived, suggesting a source-sink dynamic, where conjugative plasmids constantly generate mobilizable and putatively non-mobilizable plasmids by gene deletion. Yet, in few cases such transitions resulted in the emergence of large clades of relaxases present only in mobilizable plasmids, suggesting successful specialization of these families in the hijacking of diverse conjugative systems. Our results shed further light on the huge plasticity of plasmids, suggest that many non-conjugative plasmids emerged recently from conjugative elements and allowed to quantify how changes in plasmid mobility shape the variation of their gene repertoires.

## Introduction

Bacteria acquire DNA from other cells, eventually of different species, by multiple mechanisms of horizontal gene transfer (HGT). These genes can be integrated into the genome and expressed, thus providing adaptive phenotypic shifts (de la Cruz, Davies 2000; Soucy et al. 2015). In many species, HGT is in large part, or entirely, driven by the transfer of mobile genetic elements (MGEs) carrying accessory genes of adaptive value (Rankin et al. 2011; Hall et al. 2017; Partridge et al. 2018). Some of these, plasmids and Integrative Conjugative Elements (ICE), encode a mating pair formation (MPF) machinery that includes a type 4 secretion system (T4SS) which transfers one DNA strand from the donor to the recipient cell (de la Cruz et al. 2010). In plasmids, conjugation starts by the action of a relaxase, a multi-domain protein with transesterification activity, that nicks the DNA molecule at the origin of transfer (*oriT*) and links covalently to a single strand (Gonzalez-Perez et al. 2007). The nucleoprotein filament is then presented to the T4SS by a type 4 coupling protein and transferred to the other cell where it is circularized. Finally, the replication machineries of donor and recipient cells restore the double strand of the plasmids. The amounts of contiguous DNA that can be transferred by conjugation are without equal among the mechanisms of HGT, since entire chromosomes can be transferred in one single event (Adelberg, Pittard 1965). One other specificity of conjugation is to allow transfer across bacteria that are distantly related.

Both ICEs and conjugative plasmids conjugate MGEs that are not capable of autonomous conjugation, so called mobilizable plasmids and integrative mobilizable elements (reviewed in (Ramsay, Firth 2017)). Mobilization *in trans* occurs when the element uses a conjugative system encoded by another element for horizontal transfer. The best studied cases concern plasmids lacking a functional conjugation system and encoding an *oriT* and a relaxase. The relaxase interacts with the *oriT* and then with the T4SS encoded by another element, which is used for plasmid transfer. Here, plasmids are called conjugative when they encode a presumably complete machinery for conjugation (relaxase and MPF) and are called mobilizable when they encode a relaxase gene. It has often been observed that rates of conjugation of mobilizable plasmids are lower than those of the co-occurring conjugative plasmid (Perez-Mendoza et al. 2006 ; Blanca-Ordóñez et al. 2010; Klümper et al. 2014). Other MGEs, plasmids or elements integrated into the chromosome, only have an *oriT* (O’Brien et al. 2015; Yui Eto et al. 2021). In this case, the *oriT* interacts with a relaxase encoded *in trans* that then presents the nucleoprotein to its cognate T4SS. Since such plasmids only encode *oriT* for their mobility, and these sequences are more difficult to identify using bioinformatic tools, such plasmids are hard to class in terms of mobility using computational approaches. We will refer to plasmids lacking relaxases as pMOBless. They include all plasmids that do not encode a relaxase and for which, therefore, we ignore if they can be mobilized by conjugation. Work done a decade ago has shown that many completely sequenced genomes have plasmids and, among these plasmids, a similar number of conjugative and mobilizable plasmids is found (Smillie et al. 2010). Mobilizable plasmids tend to be smaller than conjugative plasmids, presumably because they do not need to encode the large locus associated with the MPF.

Phylogenetic and functional studies have revealed that conjugative plasmids and ICEs are very similar from the point of view of their conjugation machinery and interconversions between the two are frequent (Guglielmini et al. 2011; Johnson, Grossman 2015; Cury et al. 2018). There are eight large groups of MPF, two are specific to monoderms, incl. Archaea, and the remaining are found in diderms (Guglielmini et al. 2013). There are 9 well-known classes of relaxases that vary in their sequence, protein domains and sometimes in how they catalyze chemical reactions (Garcillan-Barcia et al. 2020), and novel putative ones are being unraveled by computational studies (Coluzzi et al. 2017). The relaxase (that defines the MOB class) is one of the few key taxonomic traits of plasmids (Garcillan-Barcia et al. 2009). Recently, the analysis of average nucleotide identity (ANI) between plasmids revealed that closely-related plasmids cluster in so-called plasmid taxonomic units (PTUs) (Redondo-Salvo et al. 2020). PTUs regroup plasmids sharing more than half of their genomes, tend to have one single type of relaxase and have a characteristic host distribution, most frequently with host ranges beyond the species barrier.

Conjugative and mobilizable elements transfer many kinds of traits, including virulence factors and symbiotic islands (Nuti et al. 1979 ; Johnson, Nolan 2009 ; Carattoli 2013). Their ability to transfer large amounts of DNA per event allows the spread of complex traits encoded in long genetic loci (Kobayashi 2018; Geng et al. 2021). The broad host range of conjugation allows transfer across bacterial Phyla and beyond, *e*.*g*. between Proteobacteria and Firmicutes (Trieu-Cuot et al. 1987) or from Proteobacteria to plants (Lacroix, Citovsky 2018). This trait may explain why conjugative elements drive the transfer of antibiotic resistance genes from distantly related bacteria to nosocomial pathogens (Pedersen et al. 2018; Che et al. 2021). But conjugation is also costly because it requires production, assembly and functioning of a large protein complex that spans multiple membranes and renders bacteria sensitive to certain phages (San Millan, Craig MacLean 2019). In this context, the added cost of having a mobilizable plasmid might be low, since the conjugative apparatus is encoded by the conjugative element. Hence, relative to conjugative elements, mobilizable elements depend on an MPF encoded *in trans* but may be less costly to the cell. The dependency of these plasmids on other conjugative elements can have some advantages. Some relaxases can be mobilized by different conjugative systems *in trans* (Garcillan-Barcia et al. 2019), allowing mobilizable plasmids to increase their frequency of transfer. Hence, traits encoded in conjugative or mobilizable elements may enjoy different patterns of genetic mobility. Such traits may also transit from one type of plasmid to the other, since there is interconversions or gene flow between conjugative and mobilizable elements (Liu et al. 2013). This can be due to translocations or fusion between replicons, scissions of plasmids, or gene deletions. To understand the evolution of conjugative elements and their impact on bacterial genomes, one thus needs to understand the evolutionary interplay between conjugative and mobilizable elements.

While the macro-evolution of the machinery for conjugation, the MPF, has been described in detail (Guglielmini et al. 2013), we know relatively little of the general patterns of evolution of plasmids relative to their categorization as mobilizable or conjugative. To shed light on this issue, we analysed a large set of plasmids in terms of their mobility by conjugation. Plasmids are better suited for such studies than ICEs, because they are better known and easier to identify accurately in complete genome sequences. To understand the evolution of conjugative and mobilizable plasmids, we assessed the relative frequency of each relaxase in mobilizable and conjugative plasmids and used the relaxases as phylogenetic markers to trace past events of change. These analyses revealed a source-sink dynamics where conjugative plasmids frequently produce mobilizable and MOBless elements. These transitions are associated with major disruptions of gene repertoires and shed light on how mobilizable and MOBless plasmids are formed. They also suggest that relaxases endure different selective pressures in conjugative and mobilizable elements.

## Material & Methods

### Plasmid Dataset

The dataset used in this study consist of 13,525 complete genomes from 2,421 species retrieved from NCBI RefSeq database of high quality complete non-redundant prokaryotic genomes (ftp://ftp.ncbi.nlm.nih.gov/genomes/refseq/, last accessed in May 2019). These genomes contain 11,805 plasmids. To avoid the misidentification of ICEs as conjugative plasmids in chromids or secondary chromosomes, we excluded from further study the 419 plasmids larger than 500 kb (Harrison et al. 2010; Smillie et al. 2010). The complete list of 11,386 plasmids can be found in Table S1.

### Detection of conjugative systems

The detection of conjugative systems was performed using MacSyFinder v.1.0.5 (Abby et al. 2014). Briefly, MacSyFinder uses HMM protein profiles and a set of rules (which are defined in MPF model files) about their presence and genetic organization to identify occurrences of given MPFs and relaxases in a genome. MacSyFinder was used with default parameters (HMMer e-value < 0.001, HMM profile alignment coverage > 50%) and was ran independently for each MPF type. To detect conjugative systems in bacterial chromosomes, we used protein HMM profiles and the MPF system definitions as described before (Cury et al. 2017).

To detect conjugative systems in plasmids, we changed the default procedure to allow for relaxase genes that are distant from the genes encoding the MPF. Hence, relaxases and T4SS were searched independently. We searched for relaxases using the same protein profiles with HMMer v.3.2.1 (e-value < 0.001, HMM profile alignment coverage > 50%)(Eddy 2011). The T4SS were searched using MacSyFinder with modified MPF system definitions: the modified definitions allow the relaxase to be absent and the MPF proteins to be at any distance from each other’s in the replicon (contrary to the 60 kb of space for the chromosome models). Replicons encoding both a relaxase and a complete MPF were considered conjugative.

Recently, a novel family of relaxases, MOB_L_, was identified in Firmicutes (Ramachandran et al. 2017). We made a protein profile for MOB_L_ with HMMER v.3.2.1 using the 817 proteins sequences described as MOB_L_ homologs in (Ramachandran et al. 2017). The search for elements of this family of relaxases revealed an important overlap with relaxases detected with the MOB_P2_ HMM profile, in agreement with previous works (Garcillan-Barcia et al. 2020), and we decided to exclude it from further analysis.

### Assessment of plasmid mobility

A plasmid was considered conjugative (pCONJ) when it encoded one or several relaxases, a VirB4, a T4CP, and a minimum number of additional MPF proteins. The latter threshold varies according to the MPF type: two proteins for MPF_FA_ and MPF_FATA_ and three for the other MPF types (B, C, F, G, I and T). These are values lower than previously used in (Cury et al. 2018) to make the assessment of a transition from pCONJ to pMOB conservative. A plasmid was considered mobilizable (pMOB) when it encoded a relaxase but lacked the conditions to be conjugative (misses T4CP, VirB4 or a sufficient number of the other components). Plasmids encoding more than 5 MPF proteins tend to be predicted as pCONJ and plasmids encoding 5 or less MPF proteins tend to be predicted as pMOB (Figure S1). To pinpoint cases where plasmids encode several genes typical of conjugative elements, but not enough to allow classing them as pCONJ, we classed mobilizable plasmids containing in addition to the relaxase, more than 5 other proteins involved in the conjugative transfer (either MPF proteins, T4CP or additional relaxases) as decayed conjugative elements (pdCONJ).

### Relaxase clustering, alignments and phylogeny

We clustered the 5,666 relaxase protein sequences using MMSeqs2 *cluster* (v. 9-d36de) with 99% identity and alignment coverage (on both query and target) thresholds (i.e. parameters -- min-seq-id 0.99 --cov-mode 0 -c 0.99) (Hauser et al. 2016). The protein sequences of the relaxases of each type were then aligned using MAFFT v.7.245 with the E-INS-I algorithm (parameters --genafpair --maxiterate 1000) (Katoh, Standley 2014). The multiple alignments were trimmed using ClipKIT with the ‘kpic-gappy’ algorithm which keeps only parsimony informative and constant sites and removed all sites above 90% gappyness (Steenwyk et al. 2020). The phylogenetic analyses were performed on the resulting multiple alignment using IQ-TREE (v.1.5.5, (Nguyen et al. 2015)) with the ultra-fast bootstrap option (1000 bootstraps) and with the best fitting model estimated *ModelFinder Plus* (-MPF) for each class of relaxases according to the BIC criterion. Trees were rooted using the midpoint function from the phangorn packages (v.2.5.5) for R.

### HMM-HMM alignments

MOB protein alignments were retrieved from (Guglielmini et al. 2011) and then transformed into HMM profiles (with ‘-add cons’ and ‘-M 50’ parameters) using hhmake from HH-suite3.0 program v.3.3.0 (Steinegger et al. 2019). The alignment of HMM profiles was performed using *hhalign* from the same suite of programs with default parameters. We took into consideration only bi-directional alignments with a probability superior to 70% and an e-value inferior to 10^−4^.

### Typing of plasmid replicons

To identify plasmid replicons, we used PlasmidFinder 2.0.1 with the 2020-07-13 database and default parameters (Carattoli et al. 2014). We identified 200 replicon groups in 4827 of the 11386 plasmids.

### wGRR estimation

We searched for significant similarity (e-value <10^−4^, identity ≥ 35%, coverage ≥ 50%) among all pairs of plasmid proteins using MMseqs2 (v. 9-d36de). The best bi-directional hits (BBH) between pairs of elements were then used to calculate the weighted gene repertoire relatedness (wGRR) (Cury et al. 2018):

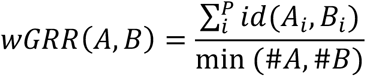

Where *Ai* and *Bi* are the *i*th BBH pair of P total pairs, *id(A*_*i*_,*B*_*i*_) is the identity between the BBH pair, and *min(#A,#B)* is the number of genes encoded by the plasmid encoding fewer genes. The wGRR varies between 0 and 1. A wGRR close to 1 means that both plasmids are highly related (all genes in an element have a very similar BBH in the other) and a wGRR of 0 means that plasmids lack homologs.

### PTU predictions

The taxonomic classifier of plasmids, COPLA, was used to assign plasmids to taxonomic units (Redondo-Salvo et al. 2021).

### Average Nucleotide Identity (ANI) calculation

Pairwise ANI scores were obtained by splitting every genome into fragments of 1.0 kb using an sliding window algorithm with a step of 200 bp. The resulting sets of fragments were compared, all against all, with a 70% identity threshold over 70% of the fragment length. Reciprocal Best Hits were selected and, if the number of selected fragments covered at least 50% of the smaller plasmid, an ANI score was assigned. The network layout was obtained using Gephi (Bastian et al. 2009).

### Plasmid ORFeome network analysis

We used AcCNET v 1.2to build the plasmid ORFeome network (Lanza et al. 2017). Homologous protein clusters were generated using kClust (v. 1.0) (Hauser et al. 2013), with >95% protein identity, >80% alignment coverage and clustering E-value < 10^−14^. All edges were assigned equal weights. The network layout was obtained using Gephi (Bastian et al. 2009).

### Inference of ancestral states

We inferred the ancestral state of each plasmid mobility type (pCONJ, pMOB and pdCONJ) with PastML (v1.9.33) (Ishikawa et al. 2019). We used the maximum-likelihood algorithm MPPA (marginal posterior probabilities approximation) and the F81 model, as recommended by the authors. The MPPA algorithm chooses for every node a subset of ancestral states that minimizes the prediction error measured by the Brier score. Hence, it may keep multiple state predictions per node but only when they have similar and high probabilities. To avoid over-estimation of the transitions between states, we counted transitions only when the ancestral node had a unique state that was different from the derived state. This may result in a conservative estimate of the total number of transitions. Out of the 287 observed transitions for all trees combined, 38 were removed because of this stringent criterion, leaving 249 transitions to be analyzed.

### Cumulative probability of transitions

For each tree, we computed the patristic distance matrix using the cophenetic.phylo function from the package ape (v5.3) for R (Paradis, Schliep 2019). Based on the patristic distance matrices, we measured the patristic distance separating each relaxase in our dataset from the closest relaxase in the phylogenetic tree that was associated with a different mobility type. For relaxases not present in the tree, i.e. not a representative of a cluster, we used the patristic distance separating the representative of the cluster and the closest relaxase in the tree having a different mobility type. This provides for each plasmid a patristic distance to the closest plasmid of a different type. The empirical cumulative distribution function of these patristic distances was computed for each MOB family and each mobility type, using the ECDF built-in function for python (v3.6.10). Empirical cumulative distribution plots were computed using, the ecdfplot function from the seaborn package (v0.11.0) for python (Waskom 2021).

### Mobile plasmid-pMOBless associations

We paired each mobile plasmid with the MOBless plasmid with the highest wGRR. To avoid the over-representation of the same transition event, if a MOBless plasmid was paired that way with multiple mobile plasmids, we only kept the mobile/MOBless pair with the highest wGRR value.

### Statistics

Unless mentioned otherwise all statistics were performed within R (v3.6.2). Statistics between two variables were done using standard non-parametric tests (Wilcoxon test). Chi-tests were performed using the R (v3.6.2) chisq-test built-in function.

## Results

### Co-occurrence and wide distribution of conjugative and mobilizable plasmids

We identified relaxases and conjugative systems among 11,386 plasmids smaller than 500kb (to avoid mis-assigned secondary chromosomes) from the completely assembled genomes of RefSeq (Table S1). Plasmids were classed in four classes (Figure 1A): 53% of the plasmids lacked relaxases (pMOBless), 23.2% were classed as conjugative (pCONJ) because they encode a relaxase and a sufficient number of MPF genes, 2.8% were classed as pdCONJ because they encoded more than 5 MPF genes but lacked the quorum of genes required to be conjugative, and 23.8% were classed as pMOB because they encoded a relaxase and 5 or less MPF genes. Hence, mobilizable and conjugative plasmids are present at similar frequencies and make up about half of all plasmids, consistent with previous results with smaller datasets (Smillie et al. 2010). The average size of plasmids varies with mobility type, with pCONJ (median size of 111.7kb) being slightly, but significantly, larger than pdCONJs (96kb) and the latter being much larger than pMOBs (37kb, P<0.001, Wilcoxon tests, Figure S7). Many plasmids encode Insertion Sequences (IS) that are known to promote genetic variation. We identified ISs in all plasmids using ISEscan v1.7.2.2 default options and found the highest densities in pdCONJ and the lowest in MOBless plasmids (Figure 1B).

**Figure 1:**
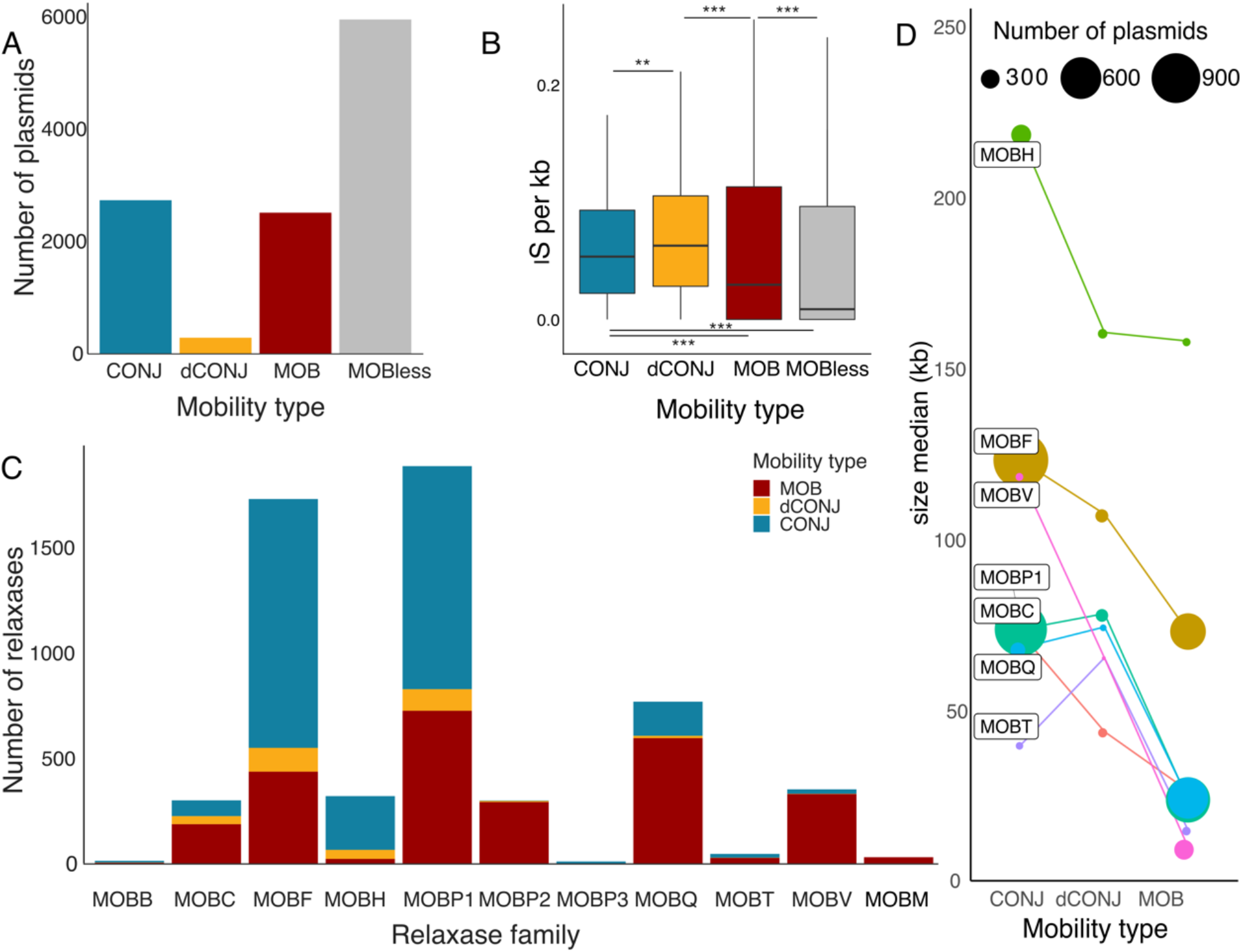
Plasmid abundance (A, C), IS density (B) and size (D) according to mobility type and relaxase class. Panel C shows the median plasmid size for each relaxase family and mobility type. The width of the circles indicates the frequency of the plasmids (see inset legend).

Our dataset of 5,666 relaxases includes all the different MOB classes, but with very different abundance (Figure 1C). By far, the most prevalent classes were MOB_F_ and MOB_P_ which account for more than 62 % of all relaxases, and are present in both pCONJs and pMOBs. It is important to note that the relative frequency of conjugative and mobilizable plasmids depends on the relaxase family (χ^2^ = 1680.6, df = 18, P < 0.001). For example, 95% of the MOB_V_ relaxases and 99% of those identified by the MOBP2 HMM profile are in mobilizable plasmids (pMOBs or pdCONJs) (Figure 1C). In contrast, 94% of MOB_H_ relaxases are in pCONJs. The average size of plasmids also varies in function of the relaxase family. Plasmids with MOB_H_ or MOB_F_ relaxases tend to be larger, whereas those with MOB_Q_, MOB_T_ and MOB_V_ are significantly smaller (Figure 1D). The trends across these types of relaxases are similar with pCONJ being in general the largest, closely followed by pdCONJ, whereas pMOB are smaller. Of note, pdCONJs to pCONJs are more similar in terms of relaxase family and plasmid size than they are of pMOB, suggesting that they were recently derived from pCONJ by gene loss.

We then assessed the distribution of plasmid mobility across bacterial taxa (Figure 2). All the taxonomic groups including more than 6 plasmids had mobilizable and/or conjugative plasmids, except the Deinococcus-Thermus. Furthermore, with the exception of the Chlorobi, which has one single plasmid, all the taxa with conjugative plasmids also have mobilizable plasmids. Hence, conjugative and mobilizable plasmids are widespread in Bacteria. The absence of these plasmids in Deinococcus-Thermus (67 plasmids) might be related with the existence of a novel transfer mechanism, called transjugation (Blesa et al. 2017). Actinobacteria and Bacilli have many more mobilizable than conjugative plasmids, which could reflect either a difficulty in identifying some conjugative systems in these clades or the previously observed over-representation of ICEs over conjugative plasmids in these clades (Guglielmini et al. 2011). If the latter is correct, then mobilizable plasmids in these clades might often be mobilized by ICEs. Alternatively, it has been suggested that non-conjugative plasmids in *Staphylococcus* spp. could be mobilized in trans by conjugative plasmids (Ramsay et al. 2016) or by transduction (Humphrey et al. 2021). Finally, given the frequency of mobilizable and conjugative plasmids, we searched to understand how often they co-occur. We found that 46.4% of the pMOBs are in genomes that encode at least one conjugative system. Hence, many pMOBs are in cells with a conjugative element and the others may eventually meet one by conjugation because most clades encode both types of plasmids.

**Figure 2:**
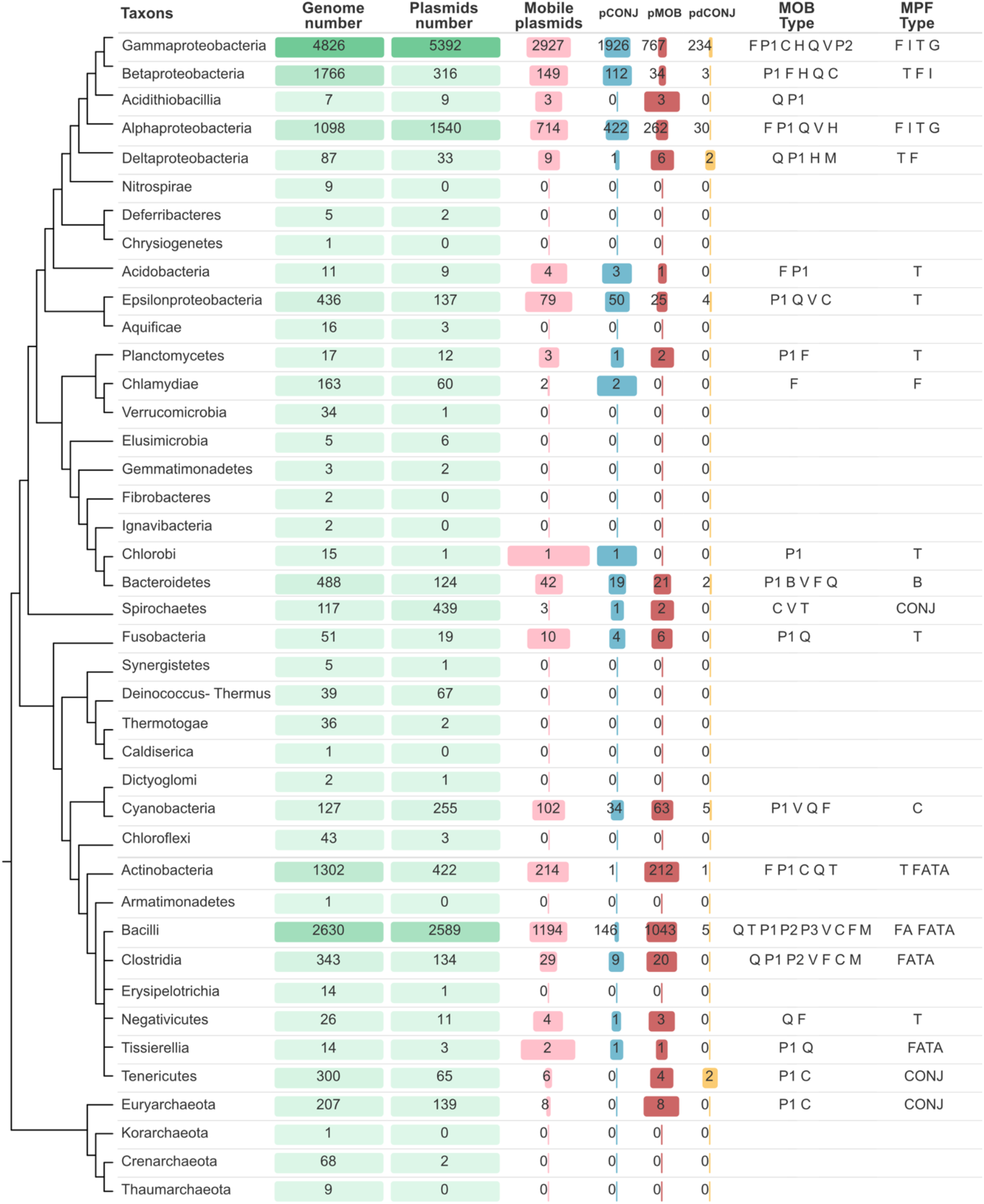
Overview of plasmids and their mobility across the tree of life. The plasmids were grouped according to their NCBI taxonomic classification of the host bacteria. This sketch tree was drawn from the compilation of different published phylogenetic analyses (Denise et al. 2019).

### The promiscuity of relaxases may favor transfer of pMOB

While relaxases from conjugative systems are expected to have co-evolved tightly with their cognate T4SS, those of pMOBs may have evolved the ability to interact with different systems to maximize their chances of transfer. To study this hypothesis, we analyzed the distribution of relaxases. Most relaxase classes are present across bacterial phyla (Figure 2). For example, the MOB_P_ class was found in Firmicutes, Proteobacteria and Bacteroidetes. In contrast, MPF types tend to be associated with specific taxa, in agreement with (Guglielmini et al. 2013), which suggests that plasmids with the same relaxase class might be transferred by different MPF types. To study the compatibility between relaxases and T4SS we studied their co-occurrence in plasmids, since it seems reasonable to assume that in most cases the relaxases and the T4SSs carried by the same plasmid can interact. When a plasmid encodes more than one relaxase, all possible MPF_TYPE_-MOB_CLASS_ combinations were taken into consideration. This analysis shows that the most frequent classes of relaxases are associated with the largest number of MPF types, but these associations are not random. For example, MOBP1 is found in plasmids with 6 different MPF types, but is preferentially associated (61.9% of the total) with MPF_T_ or MPF_I_. Similarly, MOB_F_ is mostly associated with MPF_F_ and MPF_T_ (Figure S2). These results suggest that relaxases can work in combination with a diversity of MPF types, but exhibit preferential associations with some of them.

If plasmids encode multiple relaxases to increase the range of T4SS with which they can interact, one might expect these relaxases to be from different MOB classes to increase the diversity of interactions with MPFs. Among the 11386 plasmids, only 258 encode two relaxases and 33 encode more than two. The frequencies of these co-occurrences are similar among pCONJs and pMOBs. The most frequent relaxases of pMOBs (those retrieved by MOBV/MOBP2/MOBQ HMM profiles) co-occur more frequently in pMOBs than in conjugative plasmids. In contrast, relaxase classes found in both pCONJs and pMOBs, such as MOB_F_ and MOB_P1_, or those that are mostly found in conjugative plasmids (MOB_H_) co-occur more in conjugative plasmids. This suggests that the co-occurrence of relaxases in conjugative plasmids is not often caused by fusions of pMOBs and pCONJs. Otherwise, these plasmids would have one of relaxase of each type. The co-occurrences of relaxases were as frequent within (153) as between (123) different MOB classes (Figure 3A, χ^2^ = 3.2609, df=1, P= 0.071). The MOB_V_ class is an extreme case of self-co-occurrence; it co-occurs in 24 replicons while it occurs with other relaxases in only 11 replicons. Since these results were not suggestive of maximization of the diversity of relaxases co-occurring in plasmids, we tested if they reflected the deep evolutionary relations between relaxase families using profile-profile alignments (Figure 3B). The relaxase classes MOB_F_, MOB_H_, MOB_T_, MOB_M_ and MOB_C_ showed no significant sequence homology with other classes, whereas the others were related with MOB_P1_. While fitting previous studies (Garcillan-Barcia et al. 2009), this does not show clear parallels between sequence similarity and co-occurrence in the same plasmid. Overall, these results suggest that plasmids with multiple relaxases do not seem to result systematically from the co-integration of pMOB and pCONJ neither of the optimization of the diversity of possible relaxases.

**Figure 3:**
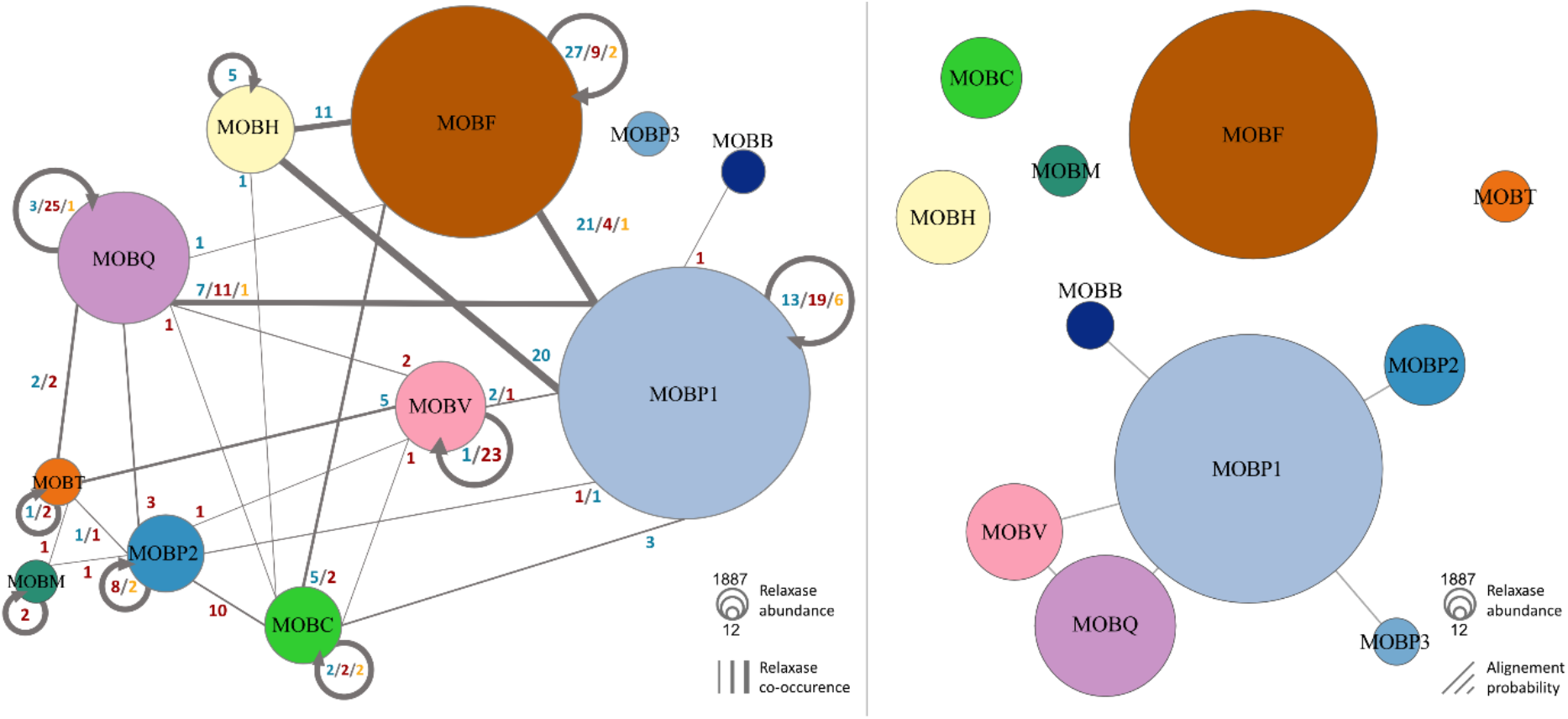
Associations between relaxases. **Panel A**. Co-occurrence (edges) of relaxases within the same plasmid. The numbers in blue, red and orange represent the number of co-occurrences found in pCONJ, pMOB and pdCONJ, respectively. **Panel B**. Representation of the results of profile-profile alignments between classes of relaxases. Edges represent bi-directional alignments between HMM profiles, and their length represents the probability of the alignment (short links represent a higher probability (up to 96%) and long links represent lower probability (down to 76%)). Are only represented alignments having a probability superior to 70% and an e-value inferior to 10^−4^.

### Plasmid size and gene repertoire evolve fast relative to the relaxase phylogeny

To start the study of the evolution of plasmid mobility we focused on the relaxases retrieved by the MOBP1 HMM profile (1,859) and MOBF (1,611) HMM profiles. MOB_P1_ and MOB_F_ are the most abundant relaxase classes, they rarely co-occur in the same plasmid, they are sufficiently conserved to be good phylogenetic markers, and they are frequent in mobilizable and conjugative plasmids. Some of the relaxases were nearly identical. To remove redundancy and accelerate computational analyses while keeping the genetic diversity of the dataset, we clustered the proteins that aligned over at least 99% of the length with at least 99% identity and picked one representative per group. This resulted in 850 MOB_P1_ and 735 MOB_F_ relaxases that were used to make phylogenetic reconstructions by maximum likelihood (see details in Methods). These proteins are representative of their clusters because most of them include only plasmids from one single mobility type (94.9% of MOB_P1_ and 92.5% of MOB_F_). The two trees are well supported in most branches. In both cases, there is a clear tendency for closely related relaxases to be in plasmids with similar MPF types. This leads to large clades over-representing one MPF type (Figure 4). In the MOB_P_ tree there is an overrepresentation of MPF_T_ that seems to suggest this was the original MPF associated with this relaxase. The MOB_F_ tree is too variable at the basal clades to make any kind of conclusion regarding the ancestral associations. Differences in plasmid size are very pronounced across the tree, since many plasmids that are close in the tree exhibit wide variations in size. Hence, plasmid size varies fast relatively to the protein sequence of the relaxases.

**Figure 4:**
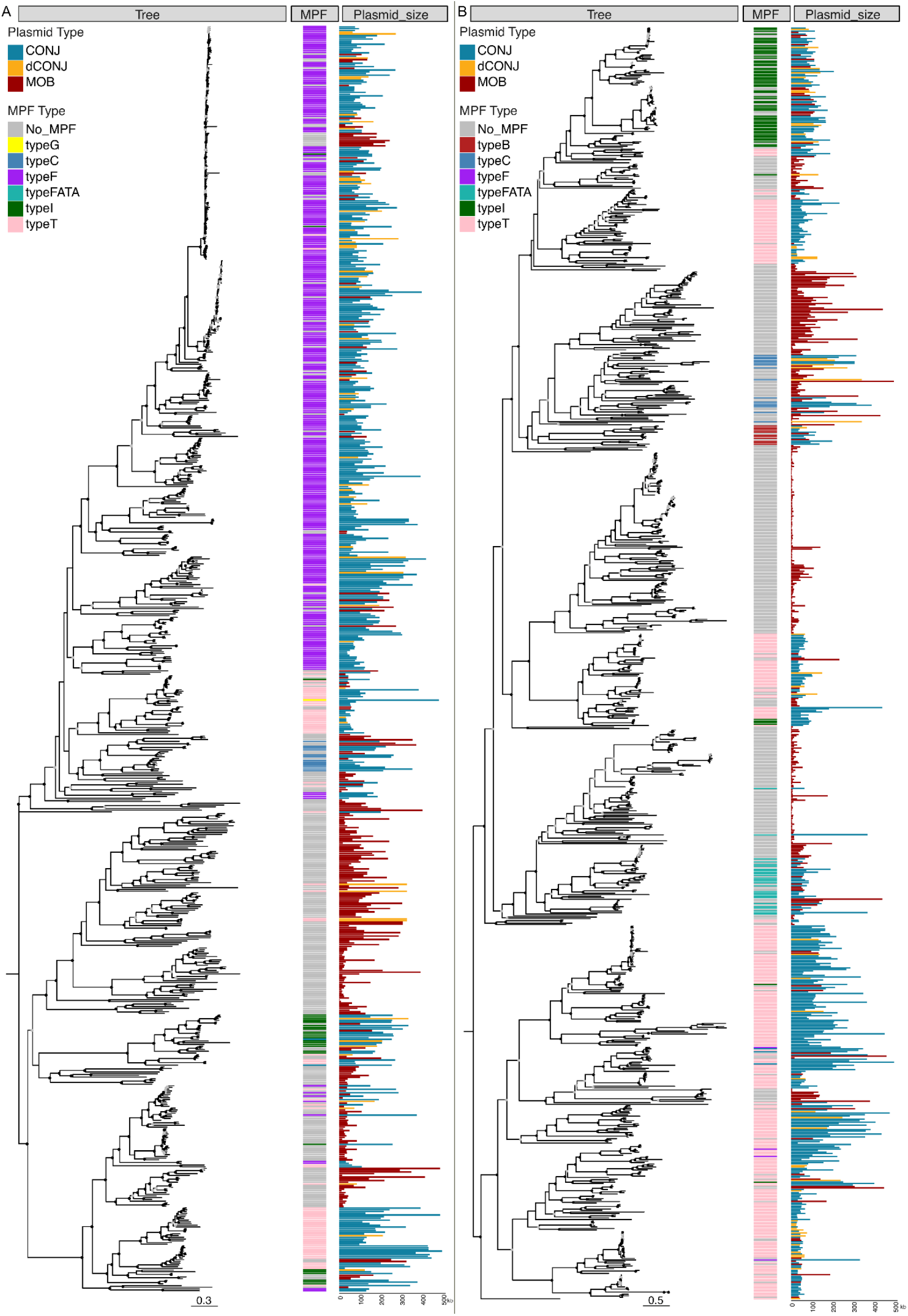
Phylogenetic trees of MOB_F_ relaxases and MOB_P1_ relaxases. Ultrafast bootstrap values superior to 75 are shown with a light grey circle and values superior to 90 with a black circle. The trees were rooted using the mid-point root. **Panel A**: The phylogenetic tree was built using 735 MOB_F_ proteins with maximum likelihood with IQTree (model WAG+F+R10 and 1000 ultrafast bootstraps (Nguyen et al. 2015)). **Panel B**: The phylogenetic tree was built using 850 MOB_P_ proteins (all retrieved with the MOBP1 HMM profile) with maximum likelihood with IQTree (model VT+F+R10 and 1000 ultrafast bootstraps).

Both phylogenetic trees divide the plasmids into large clades dominated either by pMOBs or pCONJs, but rarely containing similar numbers of both. Sister-groups are very distinctive and often separate clades of many large conjugative plasmids from those including many small mobilizable elements (although there is one group of large pMOBs in each tree). The clades with a majority of conjugative elements have many pMOBs and pdCONJ, but clades with a majority of mobilizable elements have very few pCONJ. pdCONJ are scattered across the tree, typically very close to conjugative plasmids. This is consistent with the hypothesis that such elements derived recently from conjugative plasmids by gene loss. Overall, these trees suggest early separations between sub-families of relaxases that tend to be specific for mobilizable or for conjugative elements. They also suggest more frequent conversion of conjugative elements into other type of plasmids than the other way around.

We then assessed the variations in gene repertoires for the 5,512 plasmids with the two relaxases. We calculated the weighted gene repertoire relatedness (wGRR) between each pair of plasmids, which is the number of reciprocal best hits between two plasmids weighted by their sequence similarity and divided by the number of genes of the smaller element (see Methods). A wGRR close to 1 means that plasmids are very similar (or one is a subset of the other) and a wGRR of 0 indicates no homology. Around 2500 comparisons between plasmids show very high wGRR, but most pairs of plasmids have low wGRR (Figure S3). We searched for pairs of plasmids with high wGRR but different families of relaxases. Such cases were extremely rare. We found 22 cases, of which none implicated a pair of MOB_P1_-MOB_F_ plasmids. Hence, closely related plasmids tend to have nearly identical relaxases.

Pairs of plasmids with nearly identical relaxases (≥99% identity) have similar gene repertoires (very high wGRR), except when they have different types of mobility (Figure 5, Figure S4). To understand how PTUs vary at these short evolutionary scales, we assessed their variability in clusters of plasmids having >99% identical relaxases from different MOB classes. In most cases the PTU is the same across the cluster. Yet, we found different PTUs in 18.2% of the 307 clusters analyzed, 17.5% of the MOB_P1_ and 15.6% of the MOB_F_ (Figure S5).

**Figure 5:**
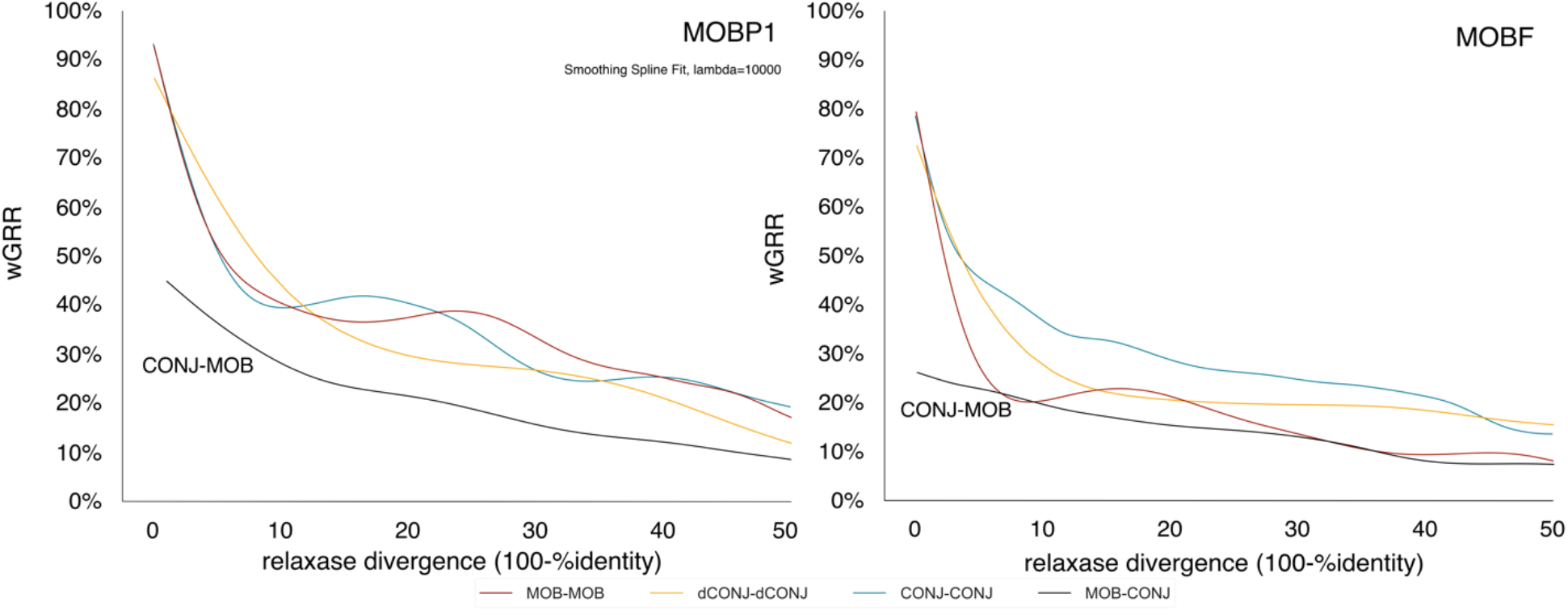
Relationship between relaxase protein sequence divergence and wGRR in pairs of plasmids classed in terms of mobility for MOB_P1_ (left) and MOB_F_ (right). Each curve represents the smoothing spline for comparisons between: pdCONJ (yellow), conjugative (blue), mobilizable (red), and comparisons between mobilizable and conjugative plasmids (black).

Diverged relaxases (<95% identity) are associated with plasmids having wGRRs lower than 50% (MOB_P1_) or 40% (MOB_F_), *i*.*e*. many such plasmids are poorly related or entirely unrelated. At 50% identity between the relaxases, the average wGRR is less than 30% (even though the plasmids have necessarily the relaxase as homolog). Interestingly, the average trends in terms of wGRR are similar for pMOBs and pCONJs, suggesting comparable rates of change in gene repertoires.

### Conjugative plasmids are the source of other plasmids

To quantify the rates of transition between mobility types, we inferred the ancestral mobility of plasmids using PastML and identified 249 terminal branches associated with a change. This analysis was made using the previously presented data of MOB_F_ and MOB_P1_ that we combined with data from the plasmids carrying other relaxases (phylogenetic trees in Figures S12-18). We focused our analysis on terminal branches because these ancestral reconstructions are more accurate and pinpoint the plasmids that have recently changed in terms mobility. The most common transitions are from pCONJ to pMOB (116, rate=0.166 Subst^-1^) or pdCONJ (77, rate=0.065 Subst^-1^) (Figure 6a). In contrast, we detected only 36 transitions from pMOB to pCONJ and even fewer from pdCONJ to the others. We found approximately similar rates of transition from dCONJ to mobilizable (0.014), vice-versa (0.013), and pdCONJ to conjugative (0.017). These values are all much smaller than the transition rates from conjugative plasmids, confirming that conjugative plasmids are the source of most of the recent transitions between plasmid mobility types.

**Figure 6:**
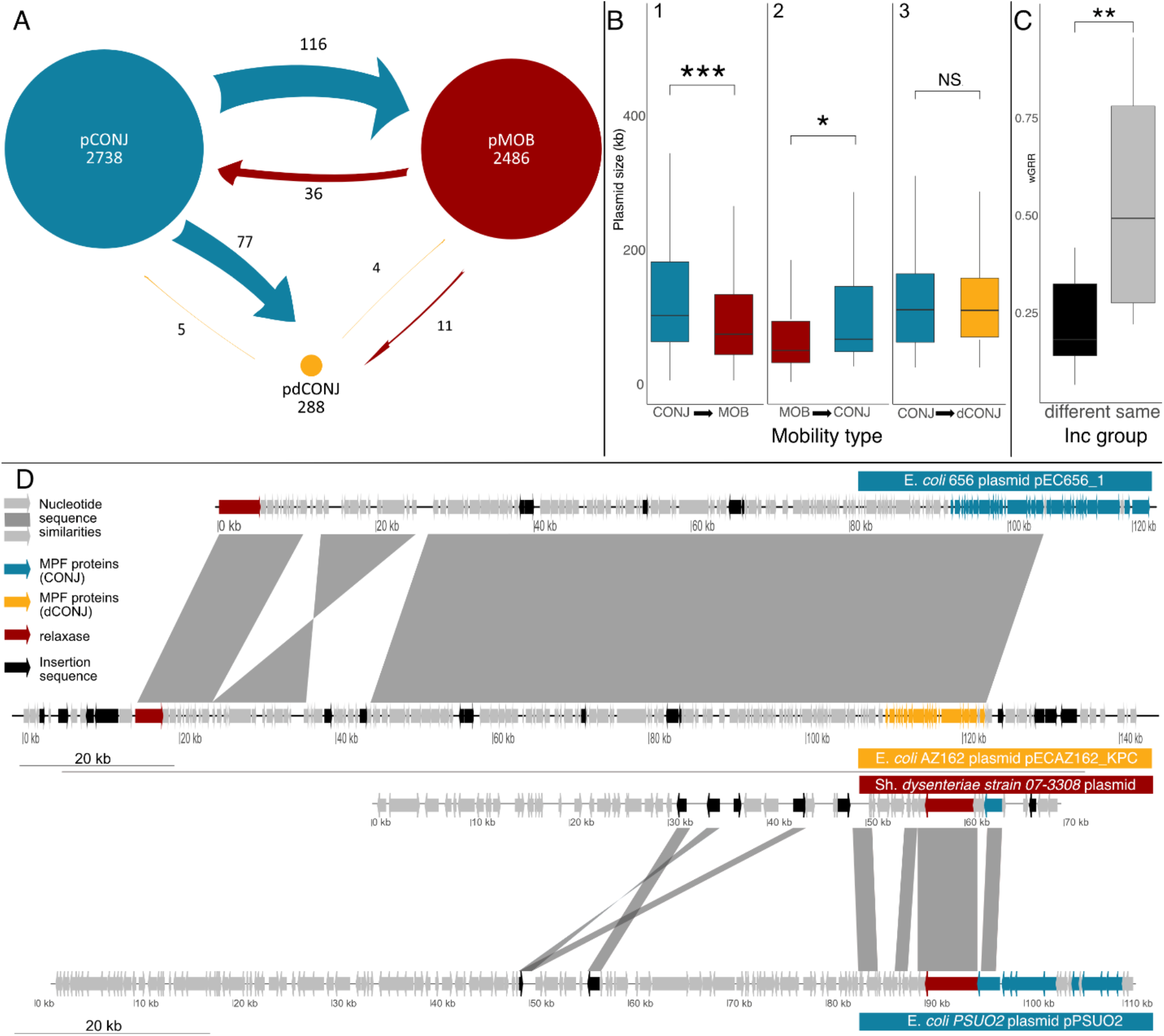
Characterization of mobility transitions. **A**. Transitions inferred in terminal branches of the relaxase phylogenetic trees. Arrows and numbers represent the direction and number of transitions, respectively. The size of circle indicates the abundance of each plasmids type. **B**. Boxplots representing the size of plasmids that recently changed in terms of mobility (in comparison to the sister-taxa plasmid in the relaxase tree having the ancestral state). Boxplot 1,2 and 3 represent the size distribution associated with transitions from pCONJ to pMOB, from pMOB to pCONJ and from pCONJ to pdCONJ respectively. **C**. Boxplot of the wGRR between pairs of plasmids where one recently changed from pCONJ to pMOB. The grey (black) boxplot represents the wGRR for plasmids of the same (different) replicon type. **D**. Comparison between selected plasmids, plotted using GenoPlotR v.0.8.10 (Guy et al. 2010), based on Blastn analysis of the plasmid nucleotide sequences (evalue < 10^−4^ and alignment length >1000 bp). Top. Comparison between a pdCONJ (yellow) and a conjugative plasmid (blue). Bottom. Comparison between a mobilizable (red) and a conjugative (blue) plasmids. For clarity, we only represent regions of homology larger than 1kb with more than 50 % identity. Genes involved in the plasmid mobility (T4SS and relaxase(s)) are highlighted. Tests of differences (Wilcoxon paired-tests): *** (P<0.001), ** (P<0.01), * (P<0.05), NS (non-significant).

The comparisons between pCONJs and pMOBs with nearly identical relaxases revealed low average wGRRs (Figure 5). This suggests that transitions between mobility types, conjugative to mobilizable or vice versa, are often associated with large changes in plasmid gene repertoires. Transitions pCONJ to pMOB can be explained by deletions of the genes encoding the conjugative system or by translocations of a relaxase from a pCONJ to a MOBless plasmid (Figure 6D). To try to distinguish between the two types of events, we computed the distribution of wGRR for pairs of closely related pMOB-pCONJ pairs. We expected to find a bi-modal distribution indicating pairs with very low wGRR originating from translocations and pairs with larger wGRR originating from gene deletions. Our analysis of MOB_F_ and MOB_P1_ failed to uncover any obvious bi-modality (Figure S6). Without excluding the possibility that translocations occur occasionally, three arguments suggest that transition by gene deletions are frequent. First, transitions pCONJ to pMOB involve a reduction in plasmid size and in wGRR (Figure 5, Figure S7, Figure 6b), but these novel pMOBs are closer in size to pdCONJ than to the typical pMOB and much larger than the average MOBless plasmid (Figure S10, Wilcoxon test, P < 0.0001). Second, the novel pMOBs often still encode a few genes of the T4SS (52% encode at least one MPF protein and 28% encode two or more), which is much more than the other pMOBs (Wilcoxon test, P <0.001, Figure S11). Third, pCONJ to pdCONJ transitions, which are presumably very recent since the conjugation locus has not yet been fully deleted in the latter, show small differences in size and wGRR (Figure 6B). This effect is not due to differences in the length of the terminal branch where we observe transitions, since those associated with transitions pCONJ to pMOB and pCONJ to pdCONJ are not significantly different (Wilcoxon test, P = 0.130, Figure S9). Instead, these results suggest that pdCONJ arise by partial deletions on the MPF locus without necessarily implicating immediate major changes in the plasmid gene repertoire (Figure 6D). These pdCONJ may then rapidly endure further gene deletions resulting in pMOBs, a process that may be facilitated by the presence of high densities of transposable elements in these plasmids (Figure 1B).

The results above suggest that changes in mobility are followed by further gene turnover that results in low wGRR between neighbor pMOB and pCONJ in the relaxase phylogenetic tree. We were particularly intrigued by the possibility that novel mobilizable plasmids could co-exist in the cell with the ancestral plasmids. If the novel pMOB maintain the replication system they had when they were conjugative, they should become incompatible with the closely related pCONJ and thus not be stably kept in the same cells. But this would affect the viability of the novel pMOB, since the closest related pCONJ are those that are most compatible with its relaxase. We therefore typed the replicons and analyzed if those that recently changed in terms of mobility also changed in terms of replication initiation proteins (Inc type). This is often the case among the few such plasmids that could be typed: 68% have an Inc type different from the one of the closest related pCONJ (17/25). The latter also have a much smaller wGRR than those that maintained the same Inc type (Figure 6C). Such changes may be part of the broader changes resulting in rapid decrease of wGRR values after transition from pCONJ to pMOB.

### Persistence of mobility type

The previous results suggest that conjugative plasmids are often the source of other plasmids. To quantify the persistence of the different types of plasmids we analyzed the phylogenetic trees of relaxases. Instead of focusing only on terminal branches of the phylogenetic trees alone, we computed the shortest patristic distances, *i*.*e*., the distances between taxa in the tree, between all plasmids of different mobility types. This allows to study plasmid lineages that are very persistent (rarely changing). We then computed the cumulative distribution functions of these patristic distances for each type of mobility (CDF) (Figure 7). A rapid initial increase in the CDF with increasing patristic distance indicates that the state is not very persistent, i.e. there are plasmids with different types of mobility at small patristic distances. In contrast, a slow increase in the CDF means that the closest plasmid with a different mobility type is usually distant in the tree. The results for MOB_P1_ and MOB_F_ are qualitatively similar and show that dCONJ plasmids are always close to other types of plasmids in the tree of relaxases. This is consistent with the frequent transitions from pCONJ and shows that these pdCONJ are transient, either because they rapidly change again their type or because they are lost from populations. Conjugative plasmids have intermediate CDFs, which is consistent with the observation that they are frequently the source of pdCONJ and pMOBs. In contrast, pMOBs exhibit a steep increase in CDF with initial increase in patristic distances, which are then followed by a much slower increase of the CDF for larger patristic distances. This is consistent with the observation that a fraction of pMOBs derived recently from pCONJ. The remaining pMOBs are placed in clades of the relaxase phylogenetic tree where there are almost only pMOBs, explaining the higher persistence of this mobility type relative to the others.

**Figure 7:**
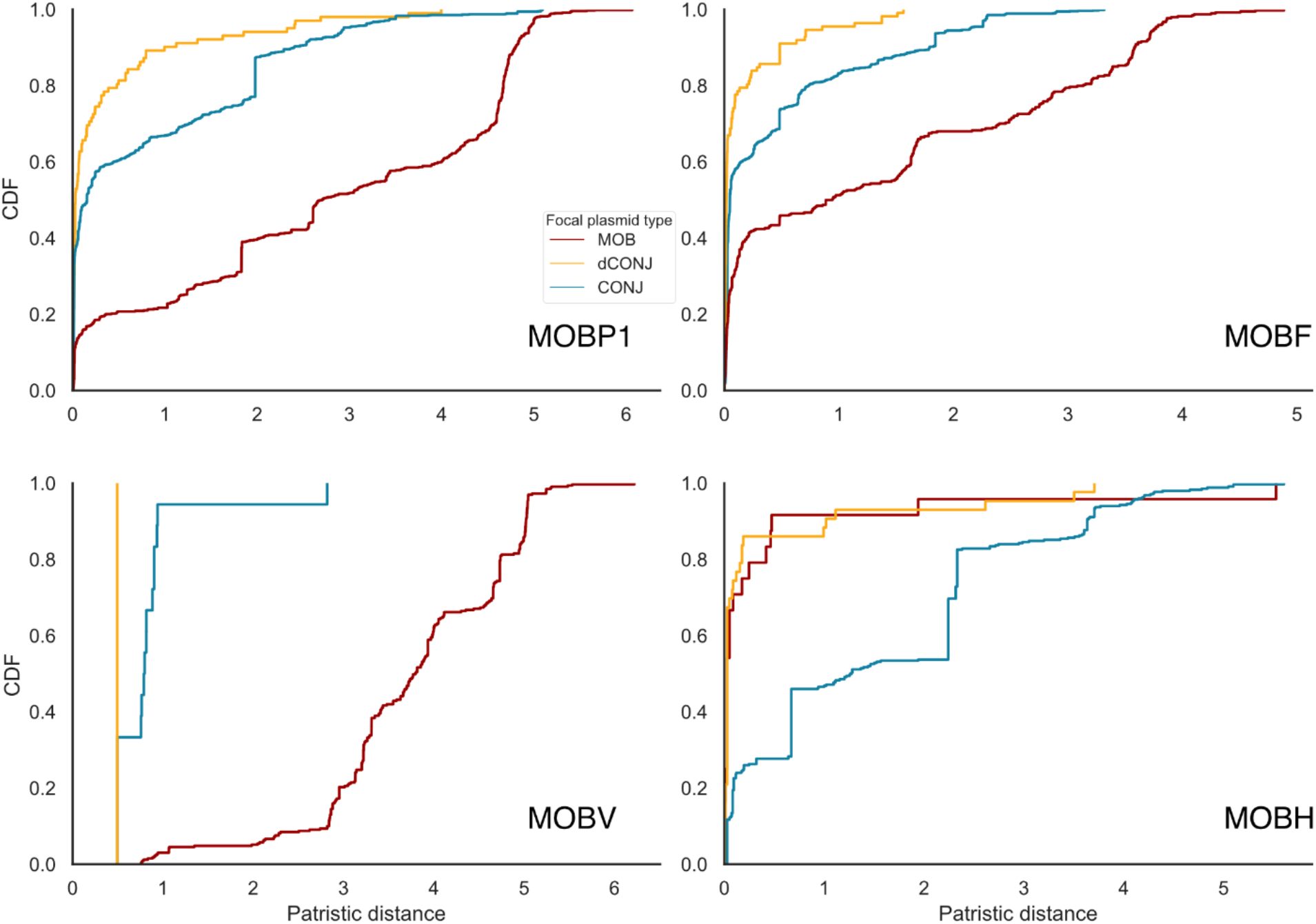
Cumulative distribution function (CDF) of the minimal patristic distances in the relaxase phylogenetic trees from a relaxase to another relaxase of a different mobility type. When the CDF approaches 1 for low patristic distances, this means that all relaxases are close to a relaxase of a plasmid with a different mobility type in the tree.

MOB_P1_ and MOB_F_ are present in many conjugative and mobilizable plasmids. To understand if the abovementioned patterns of change differ among relaxases associated with either one or the other, but not both, types of plasmid mobility, we analyzed the CDF for MOB_V_ (mostly mobilizable plasmids) and MOB_H_ (mostly conjugative plasmids). The results for MOB_V_ mirror those of MOB_P1_ and MOB_F_, with the exception that the CDF increases faster for conjugative plasmids and slower for mobilizable. This suggests that plasmids with relaxases specialized in mobilizable plasmids rarely become conjugative. In contrast, the CDF increases much slower for conjugative elements with MOB_H_, suggesting that they rarely lead to mobilizable plasmids. These results are consistent with a specialization of certain relaxases in plasmids with specific types of mobility.

### Transitions to and from MOBless plasmids are frequent

In the precedent sections we focused on conjugative and mobilizable plasmids because they can be analyzed in terms of the presence and evolution of the relaxase. Yet, around half of the plasmids lack a recognizable relaxase (MOBless, Figure 1A). If plasmids can change between conjugative and mobilizable types, they may certainly also gain or lose the relaxase. The study of the transitions to and from MOBless plasmids is difficult because they lack conserved phylogenetic markers. The only alternative to the relaxase, the replication initiation genes, are very diverse, often unrecognizable, and they can be protein or RNA genes, making deep evolutionary studies in a phylogenetic framework impossible. To study the transition between MOBless and other plasmids, we identified non-redundant pairs of MOBless-(pCONJ, pMOB, pdCONJ) plasmids that presumably diversged recently because their wGRR is high (superior to 0.75, see details in Methods). We detected 345 mobile plasmids that had one closely related MOBless plasmid: 154 pCONJs, 172 pMOBs and 19 pdCONJs (Figure 8). One might have expected an over-representation of pairs MOBless/pMOBs, since both types of plasmids tend to be small, and it only takes gain/loss of a relaxase to transition from one into the other. Yet, this is not the case and there is no significative difference between the number of pCONJ/pMOBless pairs (44.5%) and pMOB/pMOBless pairs (50%) (χ^2^= 2.8209, df = 1, P = 0.09). Hence, transitions from or to MOBless plasmids may be quite common for all plasmid mobility types.

**Figure 8:**
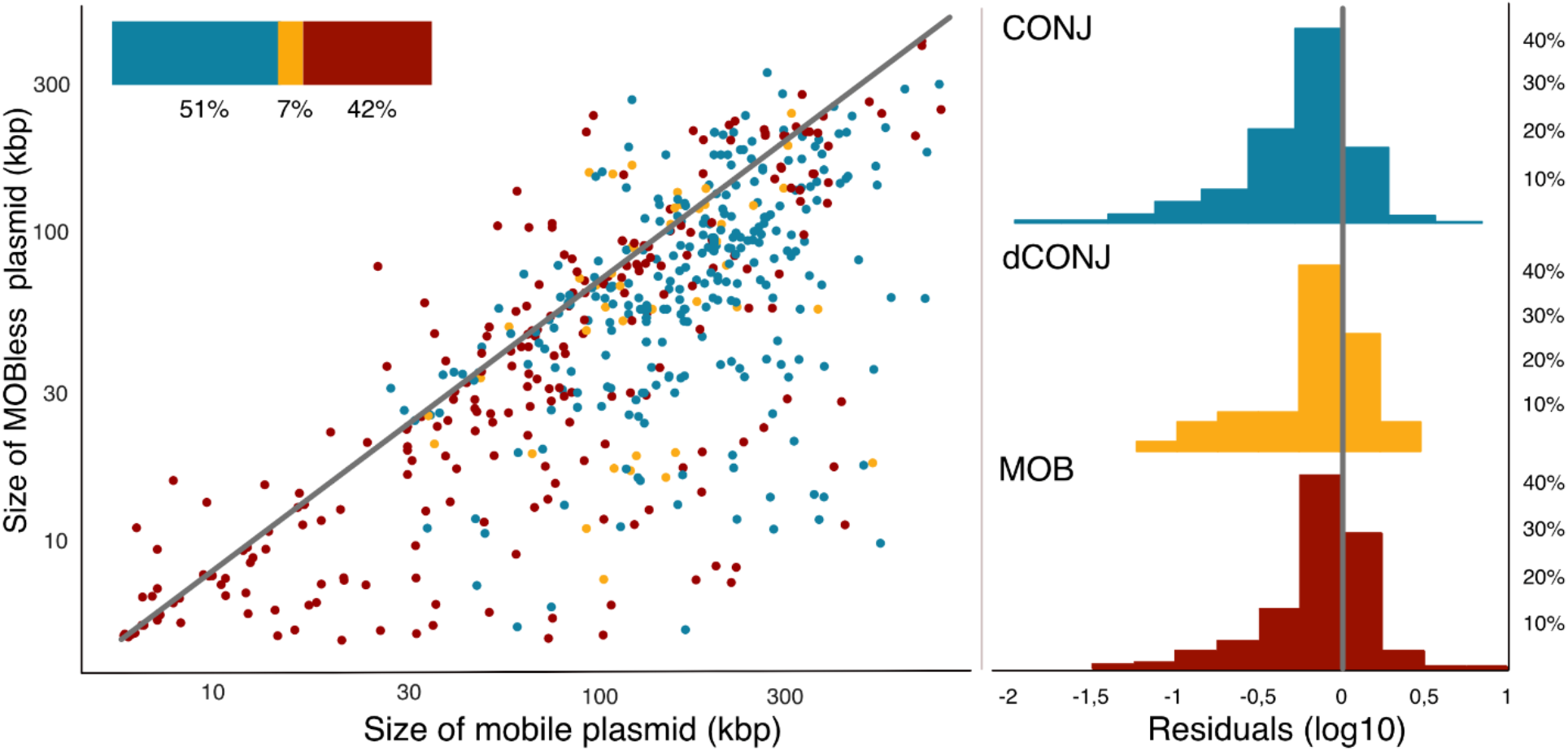
Relationship between the size of mobilizable or conjugative plasmid and the closest related MOBless plasmid. Each data point in the scatter plot represents the size of pairs of plasmids that are very similar (wGRR>0.75) and one is pMOBless whereas the other isn’t. Blue, red and yellow dots represent pairs between MOBless and CONJ, MOB and dCONJ plasmids respectively. The histograms on the right represent the distribution of the log10 of the residuals (distance to the identity line in grey). All three distributions have an average significantly lower than zero (all Wilcoxon tests, P<0.0001).

To assess the consequences of these transitions, we compared the sizes of the plasmids of each pair. When the analysis was stratified by plasmid mobility type, it revealed that MOBless plasmids are significantly smaller than the other element of the pair independently of the mobility type of the latter (Figure 8). This suggests that transitions are associated with gene losses in the novel MOBless plasmids or gene acquisition in the others (when transition occurs in the opposite sense). Alternatively, these patterns could result from co-integration of a MOBless plasmid in a plasmid with a relaxase. Yet, in this case one would expect very large differences in sizes between the pMOBless and the cointegrate. Instead, the size of pMOBless is often only slightly smaller than that of the other plasmid. This is more consistent with a model of gene deletion resulting in a transition to pMOBless.

### Conservation of PTUs when mobility changes

To understand the interplay between changes in mobility and changes in PTUs, we focused on the MOB_F_ subtree of the MOB_F12_ family of plasmids from *E. coli* and *Shigella* spp. because they are numerous and well characterized (Fernandez-Lopez et al. 2016). We analyzed these plasmids using pairwise average nucleotide identity (ANI) comparisons. This resulted in a graph where we could plot the mobility of the plasmids and the PTUs (Figure 9A). In the network one can distinguish those plasmids that have a different mobility type tend to be at the outer edge of the main PTU (PTU-F_E_) or are directly classified as different PTUs. Depending on the effect of changes in mobility on gene repertoires, they may result in conservation of the plasmid PTU, or its budding out of the original PTU, sometimes resulting in new PTUs (such as E5, F_Sh_, E41, etc.). In this specific case, the emerging PTUs seem to be associated with specific hosts (*e*.*g*. PTU-F_Sh_ to *Shigella* spp., and PTU-E5 to *E. coli* O157:H7), which may be an indication that new PTUs emerge by change of mobility and adaptation to new hosts. It should also be mentioned that relaxase genes within the members of PTU-F_E_ may have down to 90% sequence identity (example, R1 vs R100), while transitions to the new PTUs occur while their relaxases remain more than 95% identical to some members of PTU-F_E_ (examples, to PTU-E78 or PTU-E32)

**Figure 9:**
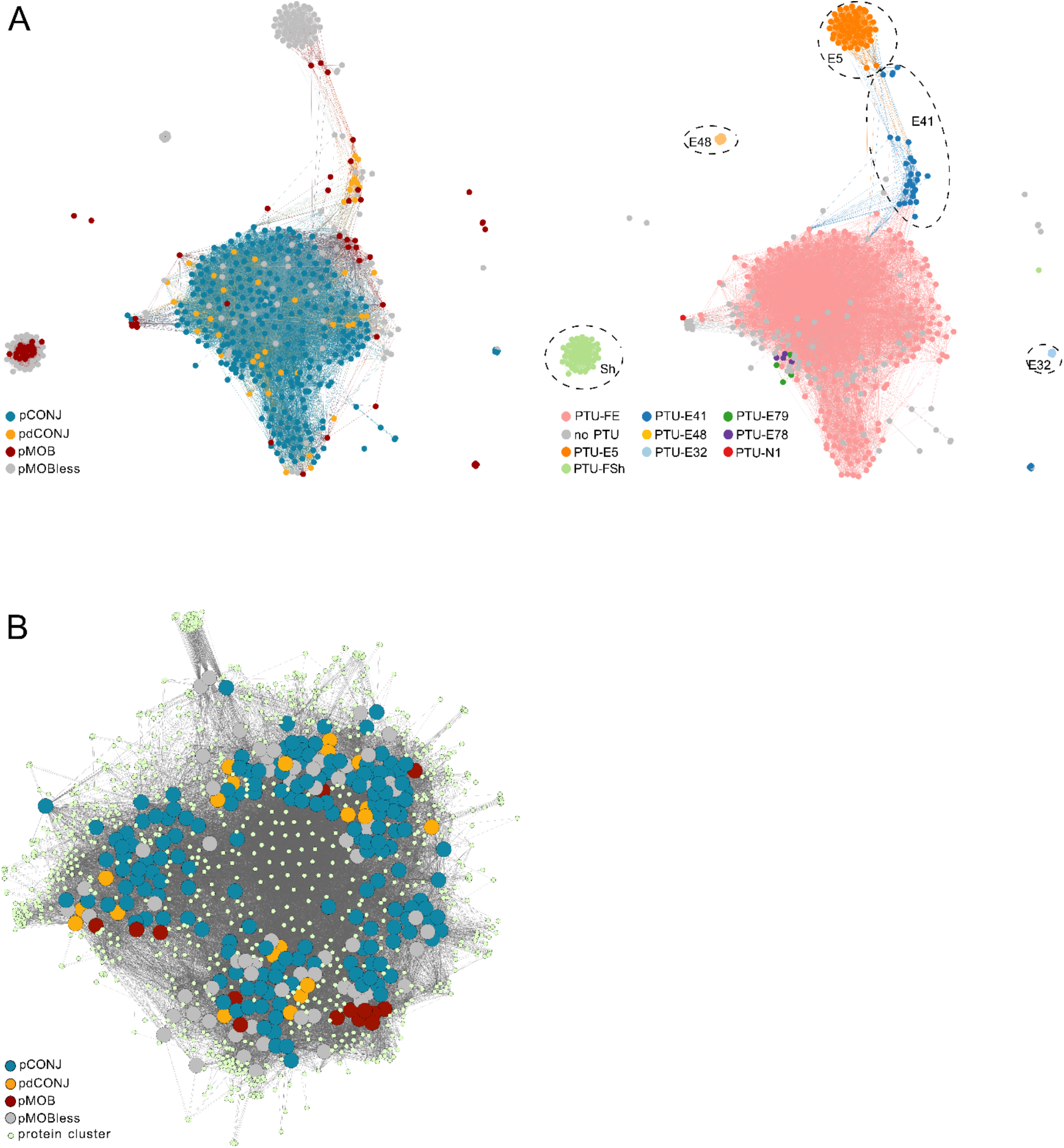
A. ANI similarity network of the MOB_F12_ plasmid family from *E. coli* and *Shigella* spp. The genomic relatedness of 353 transmissible MOB_F12_ plasmids hosted in *E. coli* and *Shigella* spp., and 140 MOBless plasmids belonging to the same PTUs was estimated by pairwise ANI calculations. The nodes correspond to plasmids and edges indicate non-null ANI scores. At the left, nodes are colored by their mobility type, and at the right, by their PTU. “no PTU” means non-assigned PTU. In the right panel, PTUs that are separated from the main group (PTU-F_E_) are surrounded by a circle. **B. ORFeome network of the PTU-F**_**E**_ **plasmids**. The proteins of 270 PTU-F_E_ plasmids were clustered at 95% identity and 80% coverage. Whenever a plasmid has a member in a protein cluster, an edge is linking them. Homologous protein clusters present in a single plasmid were removed from the figure. Plasmids are colored by their mobility type.

The largest component of the ANI graph corresponds to the PTU-F_E_ plasmids, which a PTU dominated by conjugative plasmids. We used AcCNET to analyse the ORFeome shared by the members of this PTU (Figure 9B). Results show that plasmids that changed in terms of mobility are scattered in the network. Hence, there are also many transitions of mobility type at this micro-evolutionary scale, although the changes in gene repertoires are not sufficiently large as to expel the plasmid from the PTU. Interestingly, these recent plasmid variants show sizes that are much larger than those of most mobilizable of MOBless plasmids. In this PTU, the average size of the presumably ancestral pCONJ plasmid is 117.4kb, whereas those of the novel types is very close: 118 kb for pdCONJ, 112 kb for pMOB and 95.3kb for pMOBless. This analysis at the micro-evolutionary scale is thus consistent with gene deletions being the cause of many transitions from pCONJ.

We showed above that when MOB_F12_ plasmids change in terms of mobility type they endure significant changes in gene repertoires. This means that their PTUs may also change. Indeed, we observed that PTUs of pairs of pMOBless-(pCONJ,pdCONJ,pMOB) are often different (Figure S19). When these pairs have high wGRR values (>95%), 4% of the pairs exhibit transitions between known PTUs (as shown above for some examples on PTU-F_E_). When wGRRs are lower (75-80%) these transitions increase to 9.4%. Hence, transitions in terms of mobility impact the taxonomical classification of plasmids when they implicate large changes in gene repertoires.

## Discussion

In this study we aimed at understanding the evolution of plasmid mobility. We focused on plasmids encoding relaxases because these are expected to be able to transfer horizontally. Furthermore, the relaxase is a well-conserved protein that allows the study of plasmid evolution using a rigorous phylogenetic framework, including rates and directions of change in the type of mobility. Conjugative systems, and thus relaxases, appeared early in the history of life (Guglielmini et al. 2013). Since plasmids with a relaxase can only be mobilizable if there is a conjugative system available, one may presume that conjugative plasmids arrived first and mobilizable plasmids arrived later. Since a gene deletion is sufficient for a conjugative plasmid to become mobilizable, pMOBs may have arisen very quickly after pCONJ. This fits the observation of large ancient clades of relaxases specific to mobilizable plasmids in the MOB_P_ and MOB_F_ trees. It is also consistent with the existence of MOB_P_ (specifically relaxases retrieved with the MOBP2 HMM profile) and MOB_V_ families that are distantly related with hits of the MOBP1 HMM profile and are almost exclusively found in mobilizable plasmids. Importantly, none of the relaxase classes unrelated with MOB_P_ (MOB_H_, MOB_F_, MOB_M_, MOB_C_) is specific to mobilizable plasmids. The apparent exception is MOB_T_ that is extremely rare in conjugative plasmids, but very frequent in ICEs (Soler et al. 2019). These results suggest that a few initial classes of relaxases were present in conjugative plasmids and by diversification and specialization some families became specialized in mobilizable plasmids. The existence of specialized relaxase families, manifested as specific clades in the MOB_F_ and MOB_P_ trees, suggests adaptation of these proteins to mobilizable plasmids. The alternative hypothesis, that they arose in a type of plasmid and were never transferred into another type of plasmid seems extremely unlikely given the observed rate of change of their gene repertoires, the observed transitions between types in some clades of the trees, and the previously known gene flow between plasmids (Eberhard 1990; Revilla et al. 2008; Redondo-Salvo et al. 2020).

Why would there be a specialization of relaxases? One expects tight co-evolution between the T4SS and the cognate relaxase of conjugative plasmids because they are self-mobilizable and the two components have co-evolved to interact efficiently. For example, the pCONJ RK2 and RP4 have very similar origins of transfer, but are unable to mobilize each other (Fürste et al. 1989). In contrast, mobilizable plasmids require a conjugative element to transfer and may have evolved to interact with very different conjugative systems. Accordingly, MOB_Q4_ family plasmids can be transferred by at least the MPF_I_ and MPF_T_ systems (Garcillan-Barcia et al. 2019), RSF1010 is mobilized by MPF_I_, MPF_F_ and other uncharacterized systems (Meyer 2009), MOB_P5_ mobilizable plasmids (ColE1) are mobilized by MPF_F_ and MPF_I_ plasmids (Cabezón et al. 1997), MOB_V1_ mobilizable plasmids, such as pMV158, are mobilized by MPF_T_ and MPF_FATA_ (Lorenzo-Díaz et al. 2014), and SGI1 genomic islands from *Salmonella enterica* can be mobilized by IncA and IncC plasmids (Szabó et al. 2021). A recent study showed that IncQ plasmids, which are pMOB, have exceptionally large host range (Stalder et al. 2019). Finally, while the pCONJ R388 and RP4 cannot mobilize each other, both can mobilize the unrelated pMOBs RSF1010 and ColE1 (Cabezón et al. 1994). The flexibility of pMOB relaxases may result in less efficient interactions between specific T4SS (Sastre et al. 1998). Hence, relaxases of mobilizable plasmids would evolve to interact with many different T4SS at the cost of interacting less efficiently with any single one.

Our analysis showed that gene repertoires change very fast, relative to the evolution of relaxases. The pace of change is further accelerated when there is a change in mobility type, raising the question of the genetic mechanisms driving these transitions. Transitions between types of mobility can be due to accretion/deletion of genetic material or translocation of a relaxase to another replicon. It should be noted that given the rapid pace of gene repertoire evolution, it is intrinsically difficult to distinguish large translocations from plasmid co-integrations. Still, several arguments suggest that most transitions of pCONJ to other types of plasmids tend to result from deletion of genetic material. First, the relaxase gene is not mobile in itself and the homology between closely related plasmids of different type usually extends beyond the relaxase. Second, the wGRR is rarely close to zero as expected if the putative translocation affected only the relaxase. Third, many conjugative systems become pdCONJ, and this seems clearly the result of small deletions. It is possible that many transitions between conjugative and pMOB have rapidly passed by a pdCONJ intermediate state that we were no longer able to observe in many cases. Accordingly, pdCONJ and novel pMOB are much larger than the other pMOB and closer to the size of conjugative plasmids.

The high rates of transition from conjugative to mobilizable plasmids only requires gene loss, which is frequent in bacteria (Mira et al. 2001). Deletions are stimulated by transposable elements (Cerveau et al. 2011), which we found to be more abundant in pCONJ and especially pdCONJ elements. Hence, conjugative plasmids are more likely sources of other plasmids than the converse. Selection may impact the success of gene deletions (Koskiniemi et al. 2012). Both phylogenetic trees and the analyses of patristic distances between all pairs of plasmids reveal that many transitions from pCONJ are very recent (Figures 4 and 7), suggesting that they are frequently purged by natural selection in the long term. This is indication of a source-sink dynamics, where a pool of conjugative elements (source) is constantly generating variants that require a T4SS *in trans* that tend to rapidly disappear from populations (sink). Similar processes may explain how plasmids with relaxases become pMOBless.

Some transitions from conjugative to mobilizable plasmids have given rise to specialized relaxases that may be advantageous in certain circumstances, as described above. Since most bacterial clades have both conjugative and mobilizable plasmids and half of the genomes with a mobilizable plasmid also contain a conjugative plasmid, there is plenty of opportunity for the novel mobilizable plasmid to meet a conjugative element and transfer to other cells. In such a context, why should most of the novel pMOBs or pdCONJs be purged by natural selection as suggested by our data? If the relaxase is initially specialized in a T4SS (the one previously encoded in *cis*), then the novel mobilizable plasmid may have low transfer rates because it requires a very similar T4SS to transfer. This difficulty is amplified by plasmid incompatibility. If the novel mobilizable plasmid keeps its replicon group, then it is incompatible with the most closely related pCONJ. Incompatibility means the two plasmids cannot co-exist in a stable manner and this may decrease the rate of horizontal transmission of the mobilizable plasmid to a point where it will become extinct (if not adaptive to the host). This may explain why observable transitions between conjugative and mobilizable plasmids always involve significant changes in wGRR: plasmids changing the replication machinery have a higher likelihood of surviving the initial stages after the transition. This fits our observation that many novel pMOBs have acquired a different *rep* type (have a different incompatibility family) and concomitantly endured a drastic change in gene repertoires. Other reasons may contribute to the counter-selection of novel mobilizable plasmids, including lack of coordination in expression of the relaxase and the compatible T4SS encoded in *trans*, or the existence of remnants of degraded T4SS that can be costly and even toxic to the cell.

Transitions in terms of mobility are associated with changes in the classification of replication types of plasmids and the changes in gene repertoires can also affect their classification in terms of PTUs. We show that PTUs frequently contain several mobility types within them. This is consistent with the observation that changes in mobility type are frequent in plasmid evolution, with respect to evolution of the relaxase. Many of these recent changes are not drastic enough to expel a plasmid from a PTU, as shown in figure 9B. However, plasmids that endure substantial changes in their genomes tend to separate from the original PTU. If these transitions are successful, *i*.*e*., these plasmids persist and propagate in populations, this may generate novel PTUs, as observed for E5, F_Sh_ and E41, which originated from F_E_. These new PTUs may represent specific adaptations of the plasmid genetic structure to the constraints of a new condition, in this case a novel type of mobility. Hence, substantial diversity of genomic content can be found within a PTU, but very substantial diversification of plasmids following changes in mobility often result in novel PTUs.

## Acknowledgements

Eugen Pfeifer for providing the wGRR data and Marie Touchon for providing the ISScan analysis. Manuel Arroyo for comments and suggestions. Jorge Moura de Sousa, Matthieu Haudiquet and Olaya Rendueles-Garcia for the scientific conversations. INCEPTION project [PIA/ANR-16-CONV-0005], Equipe FRM [EQU201903007835], Laboratoire d’Excellence IBEID [ANR-10-LABX-62-IBEID]. PID2020-117923GB-I00 project funded by Spanish Ministry of Science and Innovation to FdlC and MPG-B.

