## Supplemental files for "Evolution of plasmid mobility: origin and fate of non-conjugative plasmids"

**This file includes:**

Figures S1 to S19
Table S1

**Figure S1: Distribution of the MPF proteins according to the plasmid mobility types**. (page 3)

**Figure S2: Most common association between MOB families and MPF type.** (page 4)

**Figure S3: Distribution of the wGRR values between pairs of plasmids according to their mobility.** (page 5)

**Figure S4: Relationship between relaxase protein sequence identity and wGRR in pairs of plasmids classed in terms of mobility for MOBF.** (page 6)

**Figure S5: PTU transitions of plasmids within relaxase clusters.** (page 7)

**Figure S6: Relationship between relaxase protein sequence divergence and wGRR in pairs of plasmids pCONJ and pMOB for MOBP1 (top) and MOBF (bottom).** (page 8)

**Figure S7: Relationship between size variation and wGRR in pairs of plasmids that recently transited from pCONJ to pMOB.** (page 9)

**Figure S8: Distribution of size of the plasmids according to their mobility type.** (page 10)

**Figure S9: Branch length distribution for plasmids that recently transited from pCONJ to pdCONJ and from pCONJ to pMOB.** (page 11)

**Figure S10: Distribution of size of recent pMOB plasmids, pMOBs, pdCONJs and MOBless plasmids.** (page 12)

**Figure S11: Distribution of the MPF proteins encoded in pMOB plasmids.** (page 13)

**Figure S12: Phylogenetic tree of $MOB_T$ relaxases.** (page 14)

**Figure S13: Phylogenetic tree of $MOB_V$ relaxases.** (page 15)

**Figure S14: Phylogenetic tree of $MOB_H$ relaxases.** (page 16)

**Figure S15: Phylogenetic tree of $MOB_Q$ relaxases.** (page 17)

**Figure S16: Phylogenetic tree of $MOB_{P2}$ relaxases retrieved with the MOBP2 HMM profile.** (page 18)

**Figure S17: Phylogenetic tree of $MOB_B$ relaxases.** (page 19)

**Figure S18: Phylogenetic tree of $MOB_C$ relaxases.** (page 20)

**Figure S19: PTU transitions between mobile and non-mobile plasmid pairs.** (page 21)

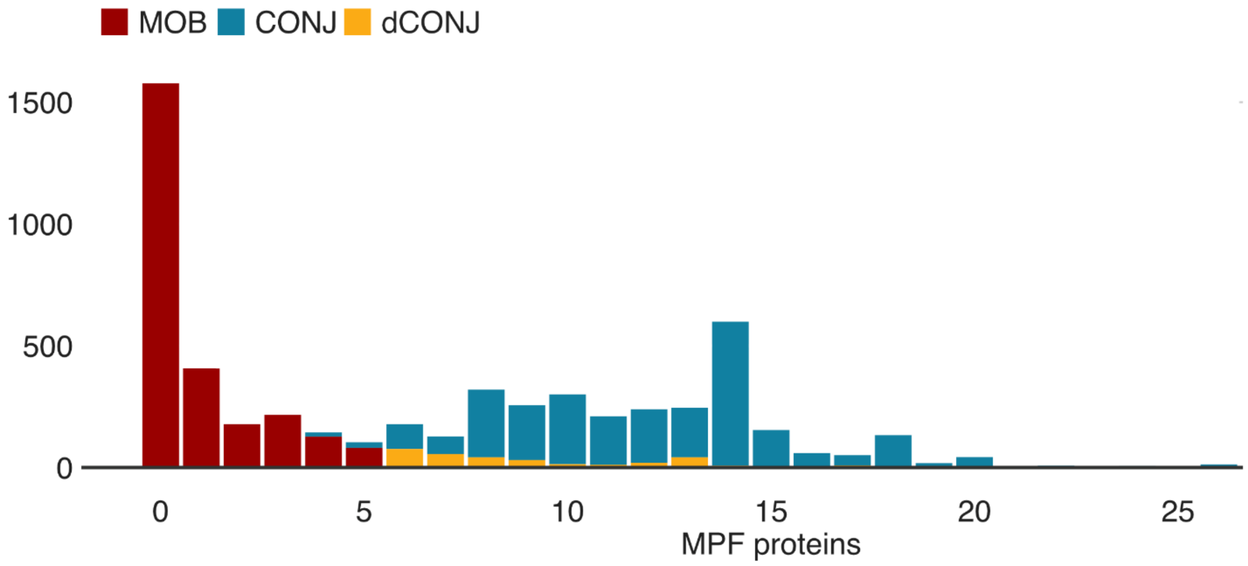


**Figure S1: Distribution of the MPF proteins according to the plasmid mobility types**. Each bar shows the number of plasmids encoding a specific number of MPF proteins according to the mobility of the plasmids.

| MOB_family | Number | MPF_associated |
|---|---|---|
| MOBP1 | 1887 | B I T C FATA FA |
| MOBF | 1731 | I T C F |
| MOBQ | 770 | G I T F FATA |
| MOBV | 354 | I C FATA FA |
| MOBH | 322 | G F |
| MOBC | 302 | T FA |
| MOBP2 | 301 | B T FATA |
| MOBT | 48 | FA |
| MOBB | 15 | B |
| MOBP3 | 12 | FATA |

**Figure S2: Most common association between MOB families and MPF type.** Most common association between MOB family and MPF type are shown for each MOB family. Associations are taken into account only if they represent more than 5% of all the association for each MPF type.

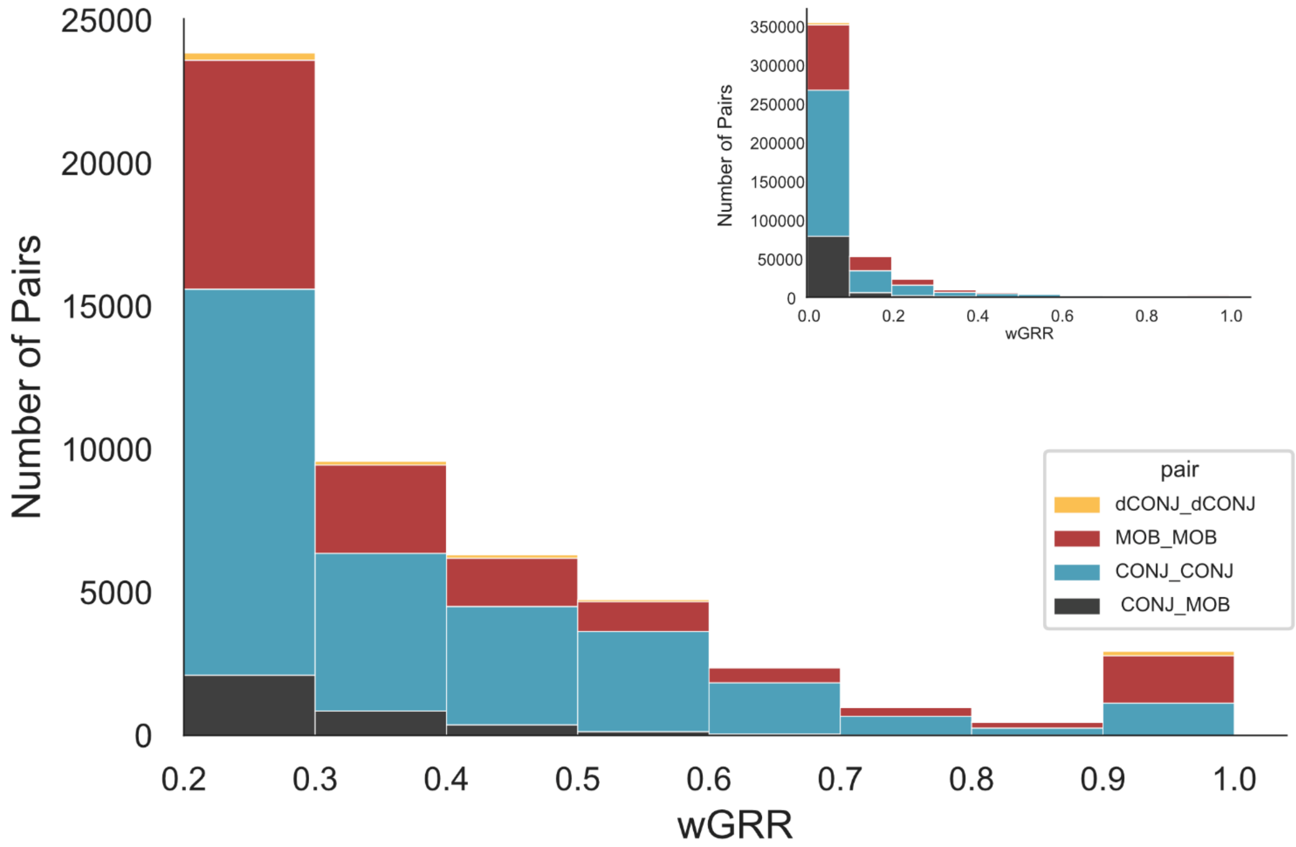


**Figure S3: Distribution of the wGRR values between pairs of plasmids according to their mobility.** For ease of readability, the distribution of wGRR values superior to 0.2 are shown in the main histogram. The complete dataset is shown in the upper right-hand corner insert.

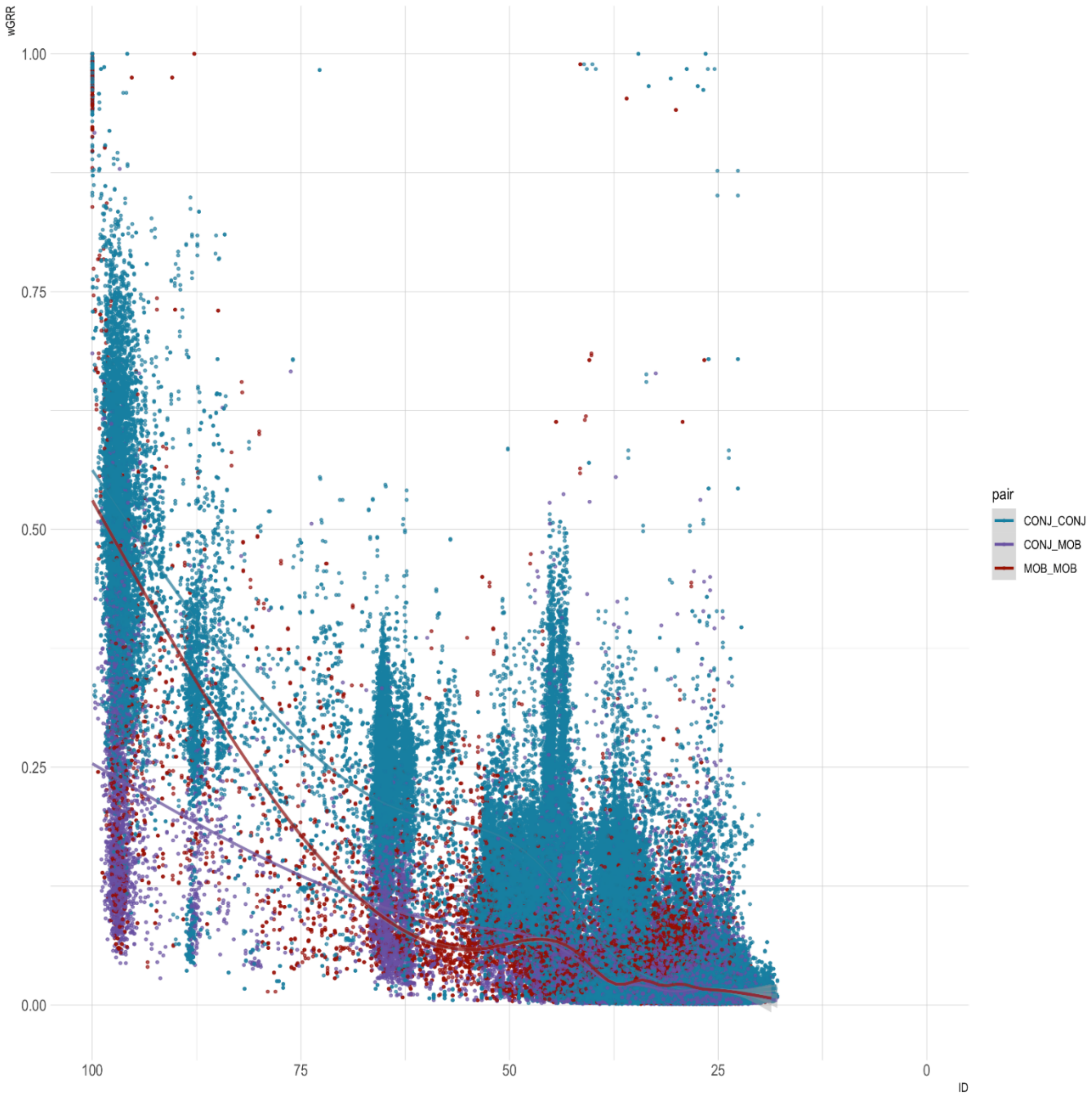


**Figure S4: Relationship between relaxase protein sequence identity and wGRR in pairs of plasmids classed in terms of mobility for MOBF.** Each point represents the wGRR between two plasmids given the identity percentage between their MOB protein. pCONJ/pCONJ pairs are represented in blue, pMOB/pMOB pairs are represented in red and, pCONJ/pMOB pairs are represented in purple.

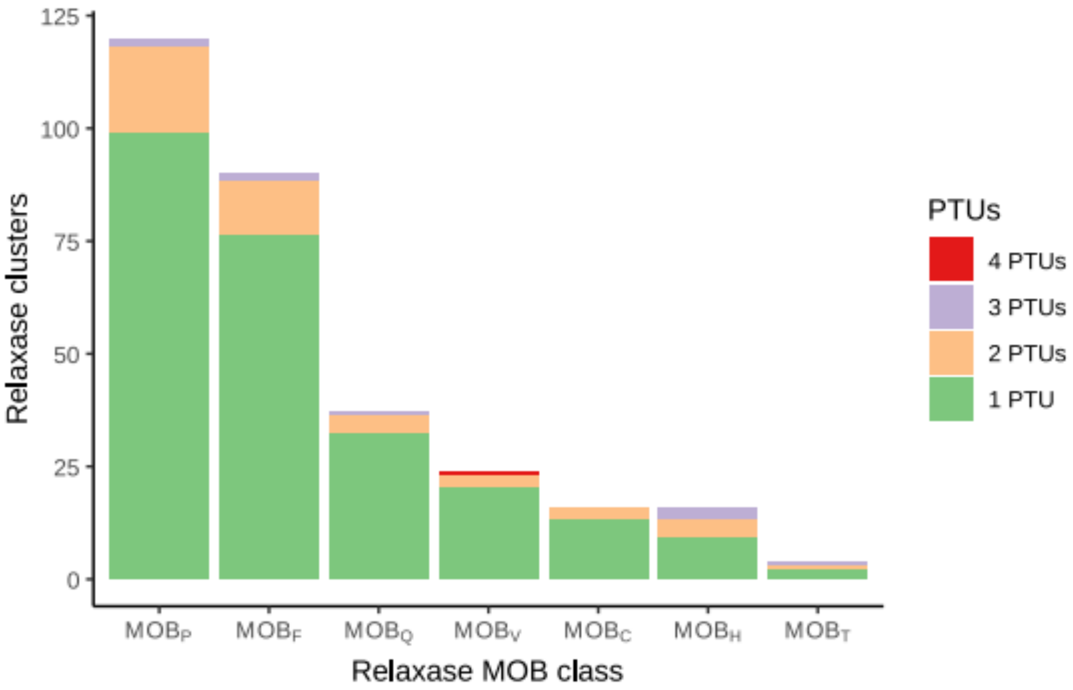


**Figure S5: PTU transitions of plasmids within relaxase clusters.** For each MOB class, clusters of >=99% identical proteins were purged from members belonging to plasmids whose PTU was not assigned. The resulting clusters containing more than one protein were inspected for the number of different PTUs they included. In blue, the clusters gathering proteins from plasmids of the same PTU (1 PTU). In orange, yellow and green, clusters whose plasmid members belonged respectively to two, three, or four different PTUs (2 PTU, 3 PTU, or 4 PTU). Number of clusters analyzed: 120 $MOB_P$, 90 $MOB_F$, 37 $MOB_Q$, 21 $MOB_V$, 16 $MOB_C$, 16 $MOB_H$, and 4 $MOB_T$.

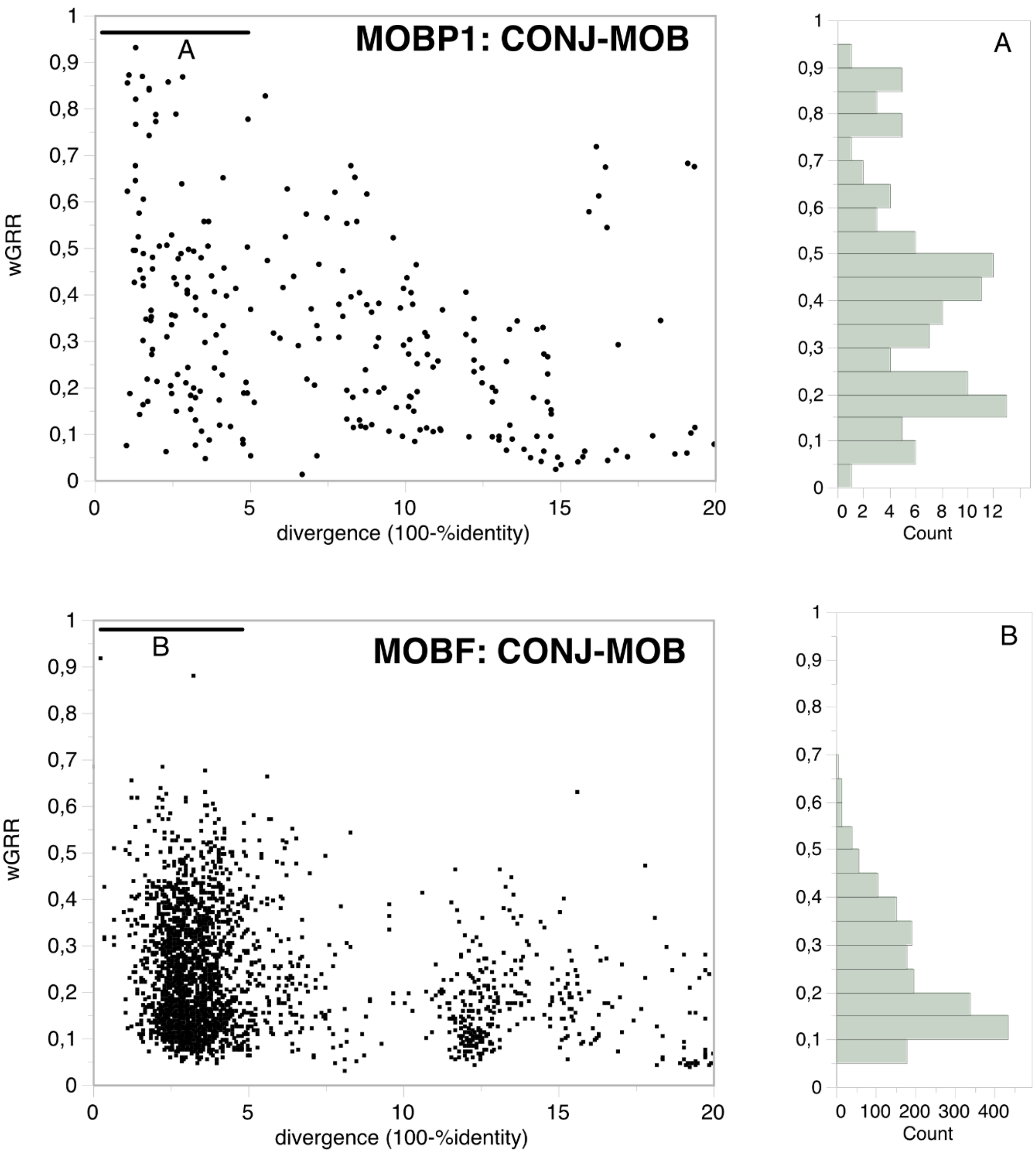


**Figure S6: Relationship between relaxase protein sequence divergence and wGRR in pairs of plasmids pCONJ and pMOB for MOBP1 (top) and MOBF (bottom).** Each data point in the scatter plots represents wGRR and the relaxase identity divergence between pairs of pCONJ and pMOB plasmids. The histograms on the right represent the distribution of plasmid pairs according to their wGRR.

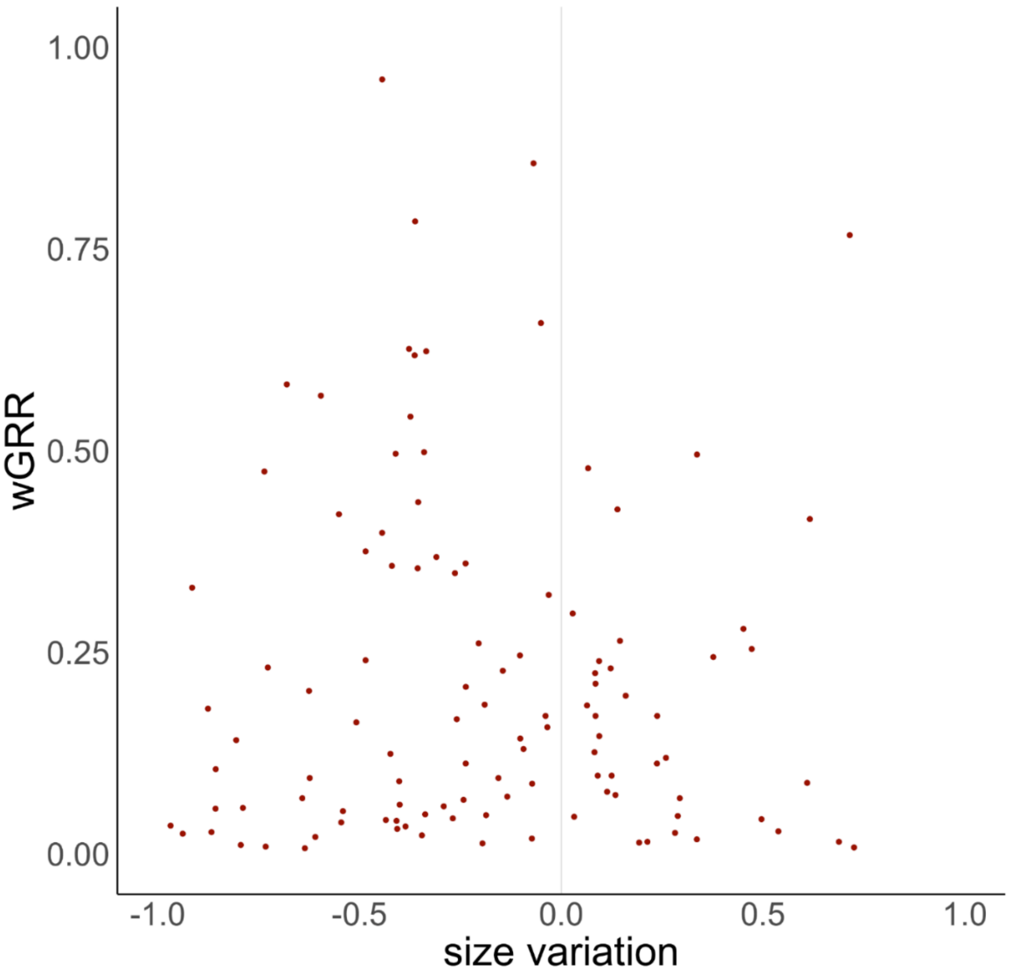


**Figure S7: Relationship between size variation and wGRR in pairs of plasmids that recently transited from pCONJ to pMOB.** Each point represents the wGRR and the size variation between each recently transited pMOB and its closet pCONJ relative. The size variation has been calculated using the formula: $(pMOB_{SIZE} – pCONJ_{SIZE})/Biggest_{SIZE}$.

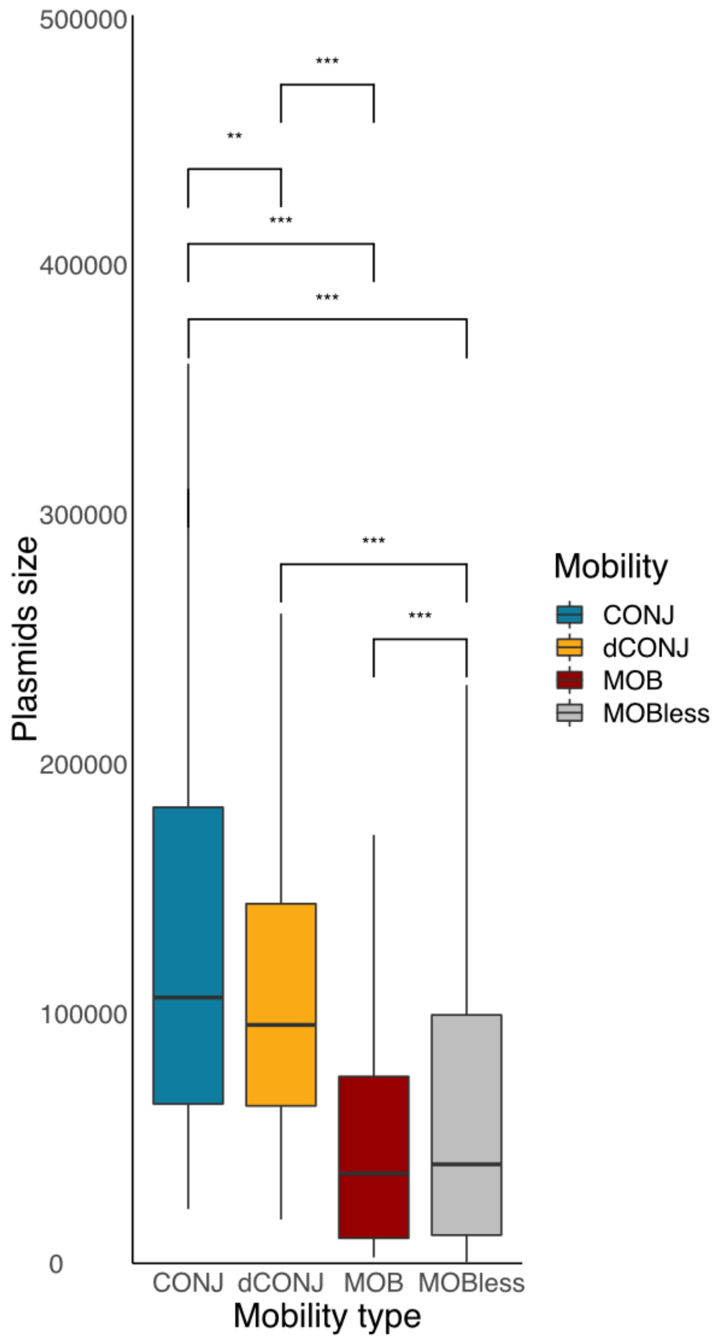


**Figure S8: Distribution of size of the plasmids according to their mobility type.** Tests of differences (Wilcoxon test): *** (P<0.001), **(P<0.01).

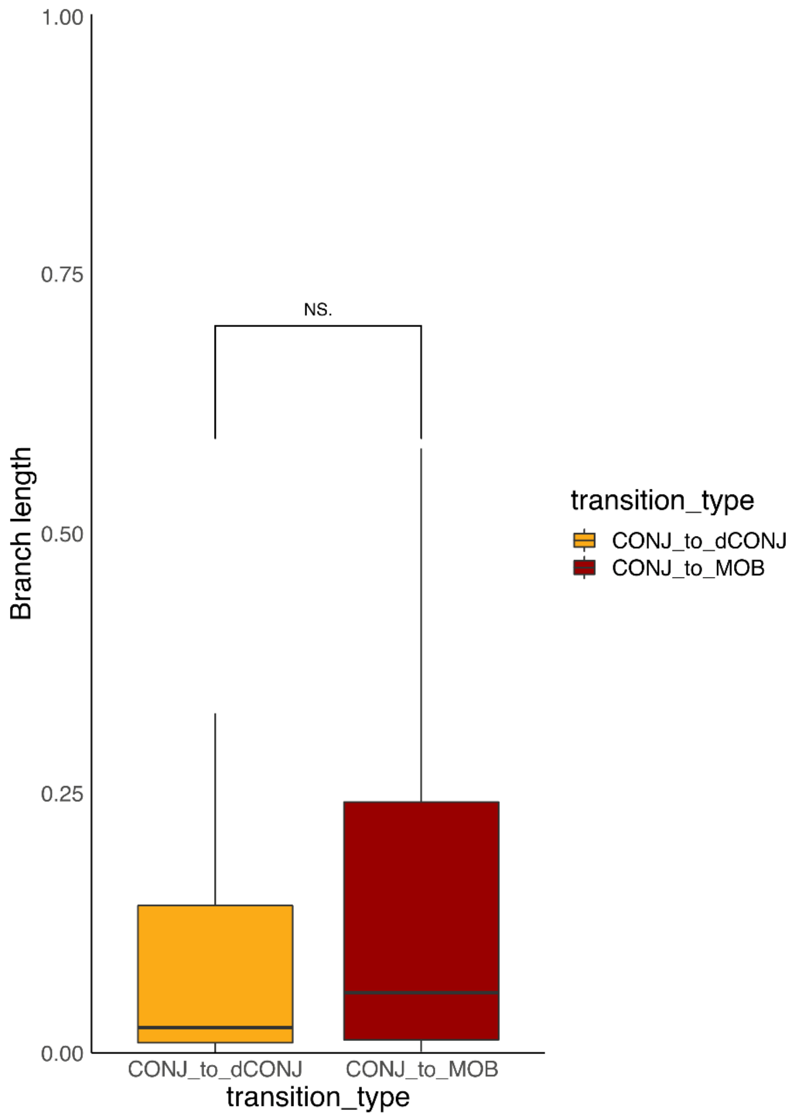


**Figure S9: Branch length distribution for plasmids that recently transited from pCONJ to pdCONJ and from pCONJ to pMOB.** Tests of differences (Wilcoxon test): NS (non-significant).

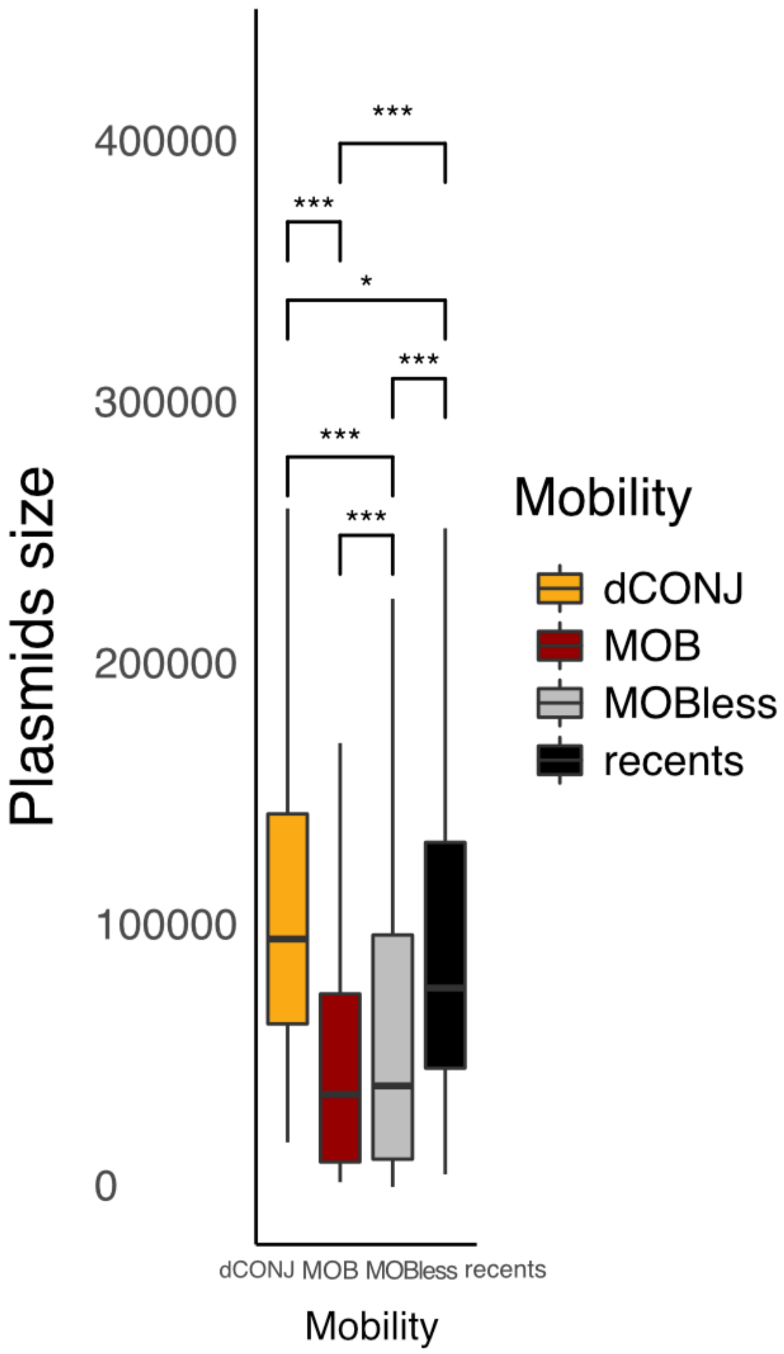


**Figure S10: Distribution of size of recent pMOB plasmids, pMOBs, pdCONJs and MOBless plasmids.** The recent plasmids are the plasmids that have recently transited (i.e. observed in terminal branches of the tree) from other types of plasmids (conjugative or dCONJ) and the remaining ones. MOBless plasmids are the plasmids without relaxase. Tests of differences (Wilcoxon test): *** (P<0.001).

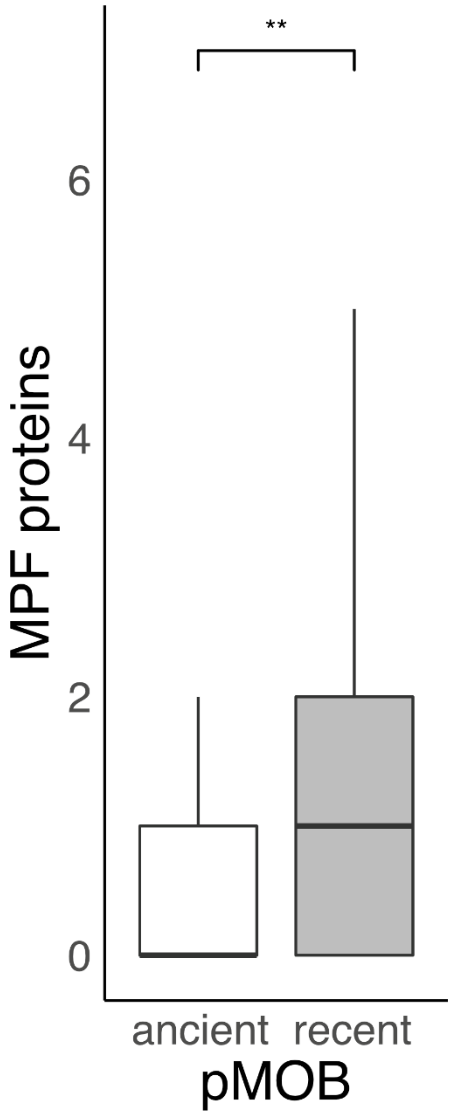


**Figure S11: Distribution of the MPF proteins encoded in pMOB plasmids.** The plasmids were split in plasmids that have recently transited (i.e. observed in terminal branches of the tree) from other types of plasmids (conjugative or dCONJ) and the remaining ones. Tests of differences (Wilcoxon test): ** (P<0.01).

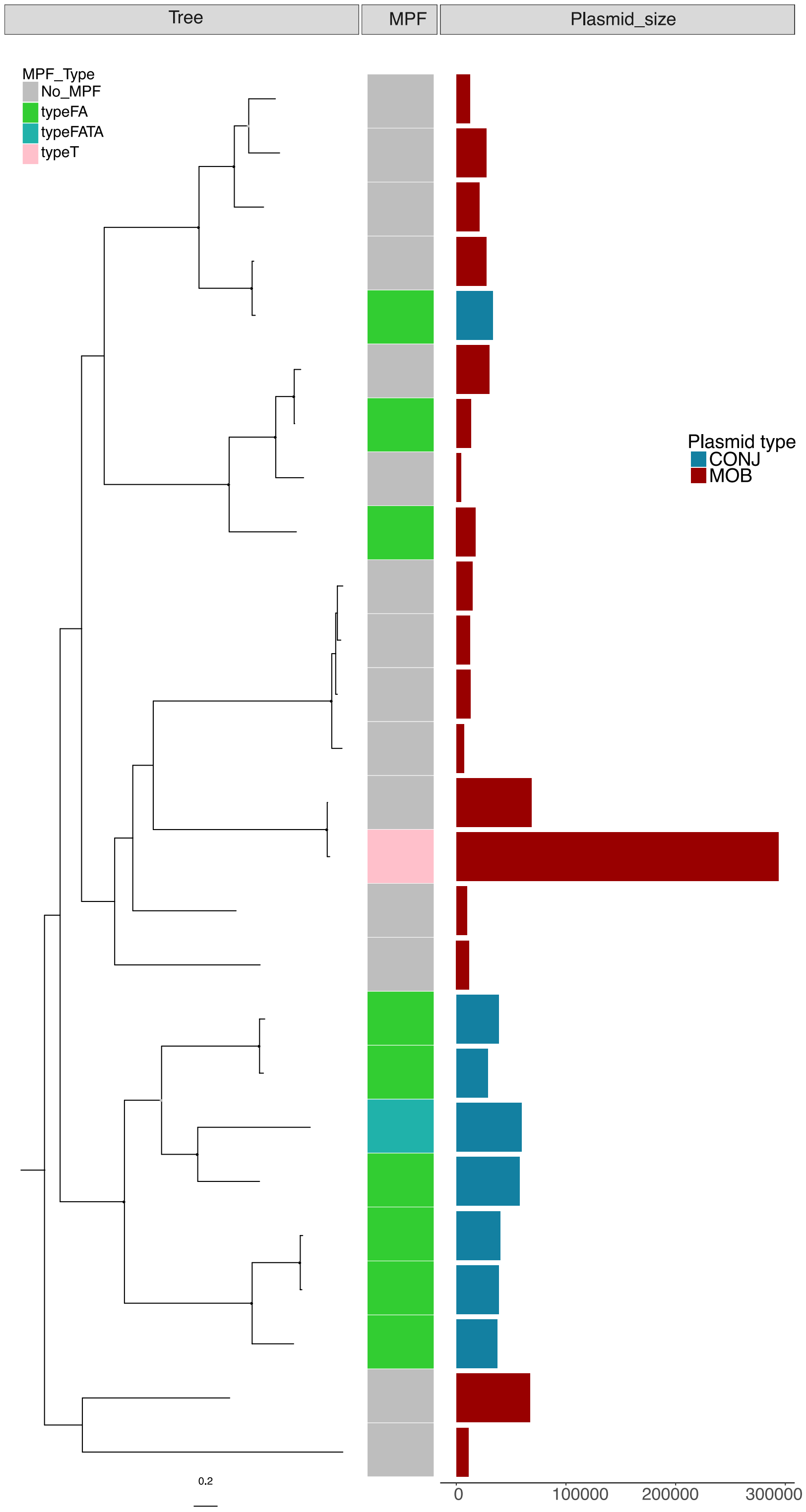


**Figure S12: Phylogenetic tree of $MOB_T$ relaxases.** Ultrafast bootstrap values superior to 75 are shown with a light grey circle and values superior to 90 with a black circle. The trees were rooted using the mid-point root. The phylogenetic tree was built using 26 MOBT proteins with maximum likelihood with IQTree (model WAG+F+R10 and 1000 ultrafast bootstraps).

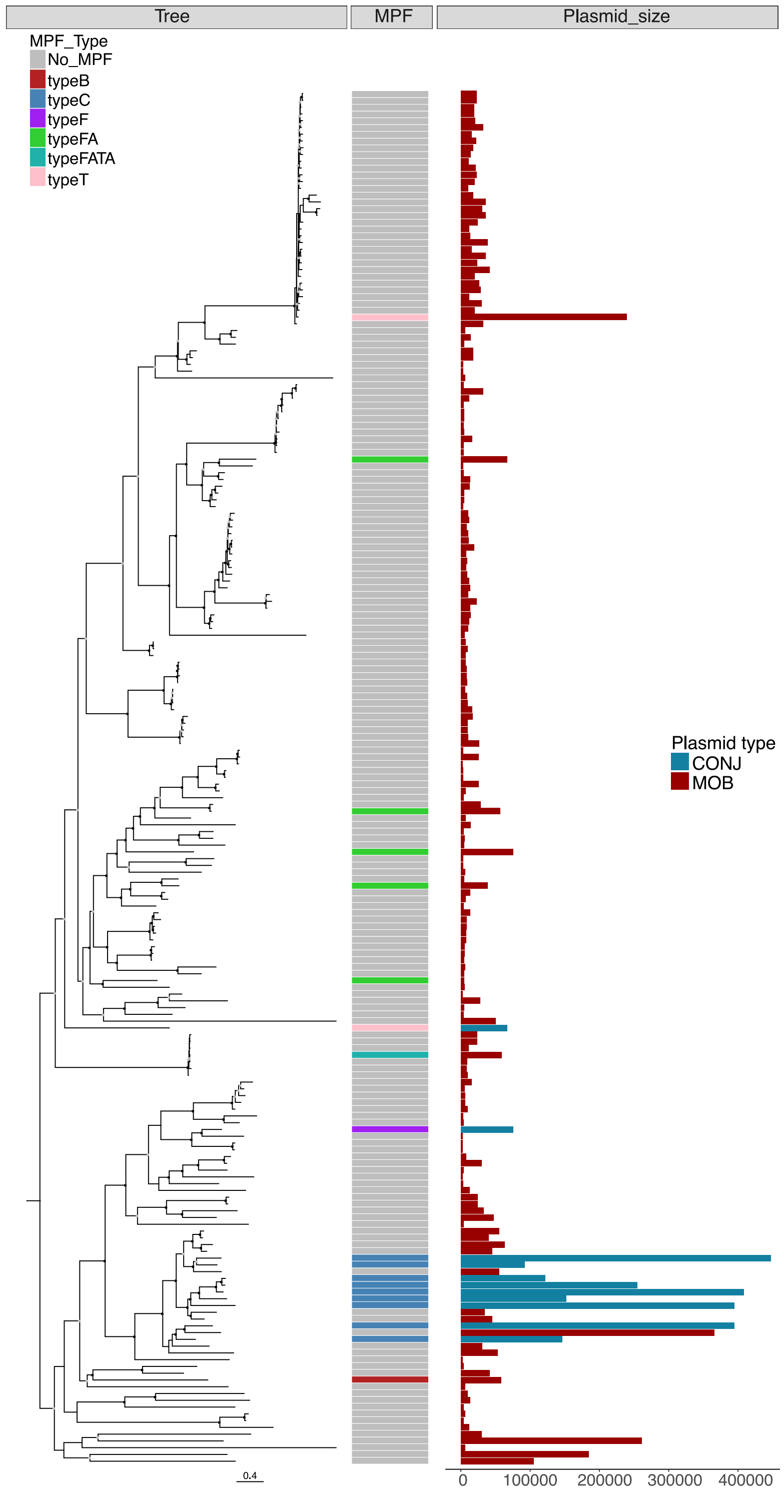


**Figure S13: Phylogenetic tree of MOB$_V$ relaxases.** Ultrafast bootstrap values superior to 75 are shown with a light grey circle and values superior to 90 with a black circle. The trees were rooted using the mid-point root. The phylogenetic tree was built using 203 MOBV proteins with maximum likelihood with IQTree (model WAG+F+R10 and 1000 ultrafast bootstraps).

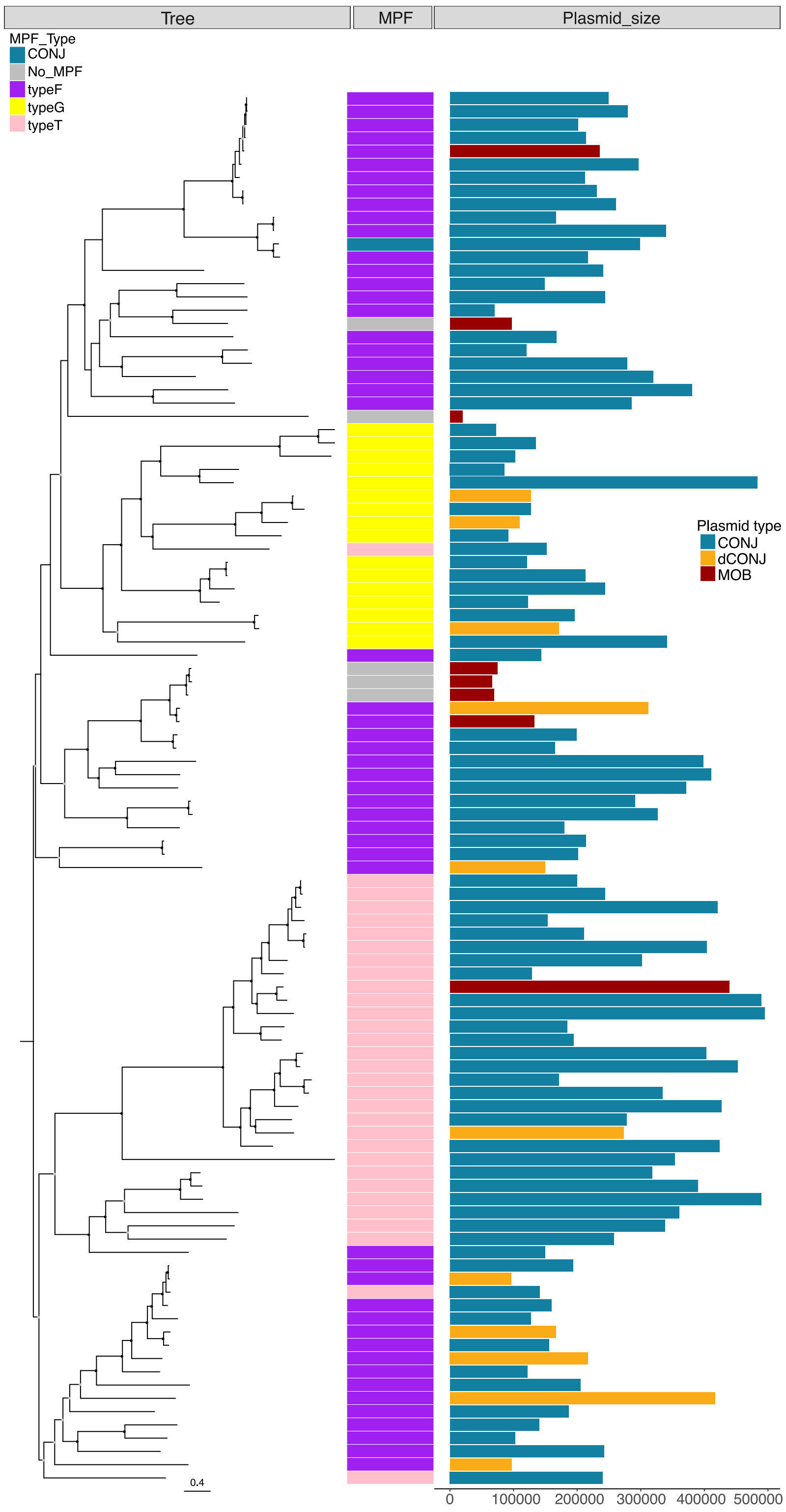


**Figure S14: Phylogenetic tree of $MOB_H$ relaxases.** Ultrafast bootstrap values superior to 75 are shown with a light grey circle and values superior to 90 with a black circle. The trees were rooted using the mid-point root. The phylogenetic tree was built using 105 MOBH proteins with maximum likelihood with IQTree (model LG+F+R7 and 1000 ultrafast bootstraps).

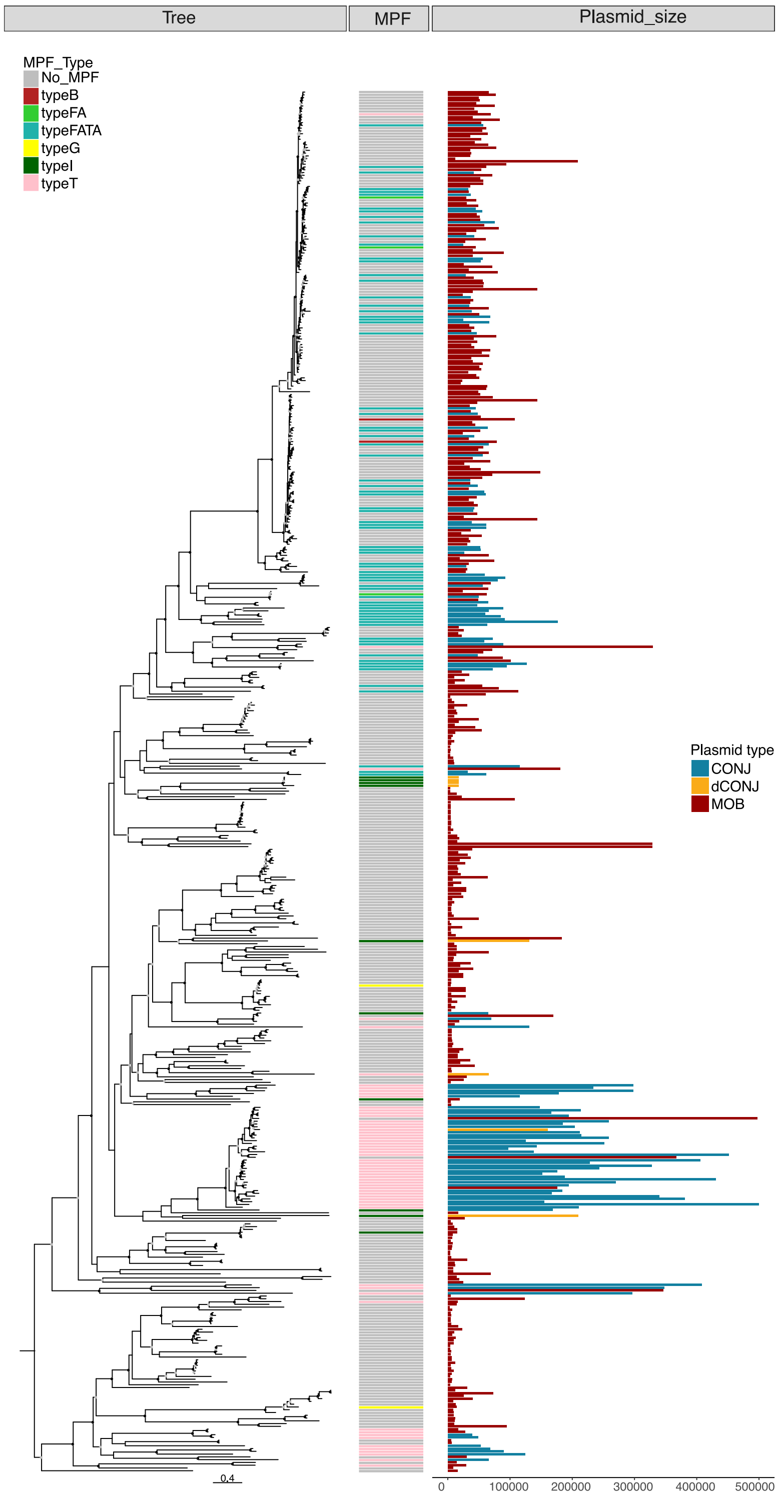


**Figure S15: Phylogenetic tree of $MOB_Q$ relaxases.** Ultrafast bootstrap values superior to 75 are shown with a light grey circle and values superior to 90 with a black circle. The trees were rooted using the mid-point root. The phylogenetic tree was built using 498 MOBV proteins with maximum likelihood with IQTree (model LG+F+R9 and 1000 ultrafast bootstraps).

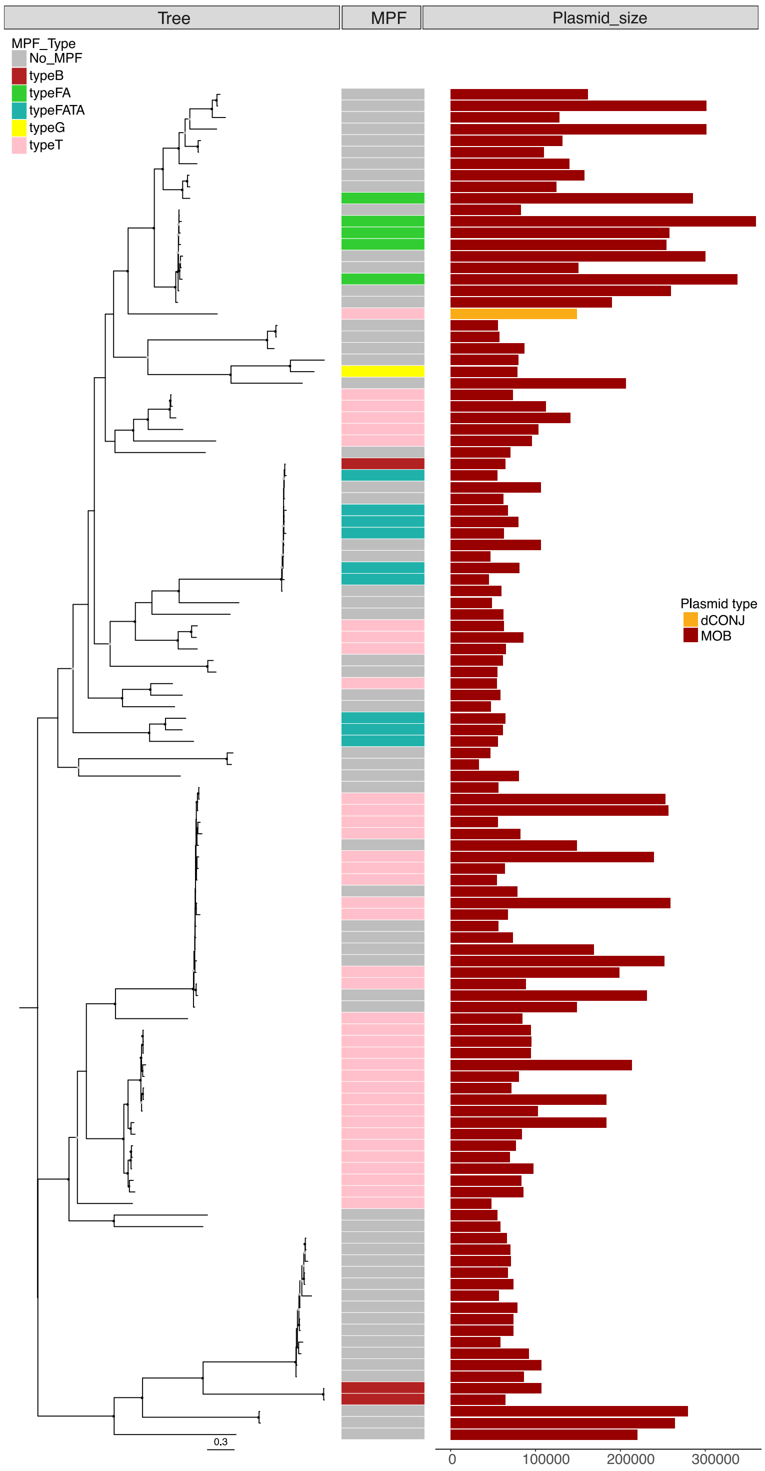


**Figure S16: Phylogenetic tree of $MOB_{P2}$ relaxases retrieved with the MOBP2 HMM profile.** Ultrafast bootstrap values superior to 75 are shown with a light grey circle and values superior to 90 with a black circle. The trees were rooted using the mid-point root. The phylogenetic tree was built using 117 MOBP2 proteins with maximum likelihood with IQTree (model WAG+F+R10 and 1000 ultrafast bootstraps).

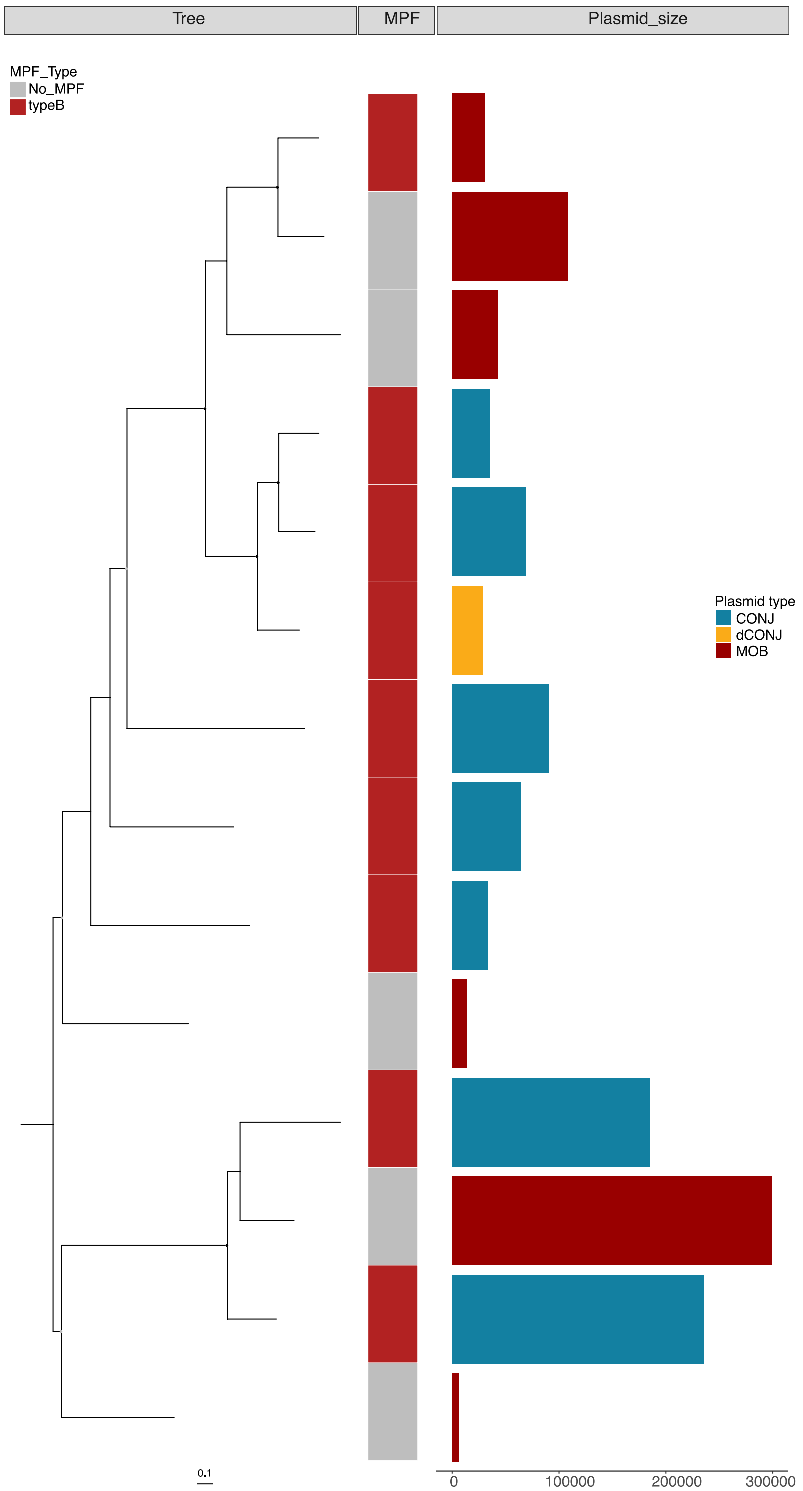


**Figure S17: Phylogenetic tree of $MOB_B$ relaxases.** Ultrafast bootstrap values superior to 75 are shown with a light grey circle and values superior to 90 with a black circle. The trees were rooted using the mid-point root. The phylogenetic tree was built using 14 MOBB proteins with maximum likelihood with IQTree (model WAG+F+R10 and 1000 ultrafast bootstraps).

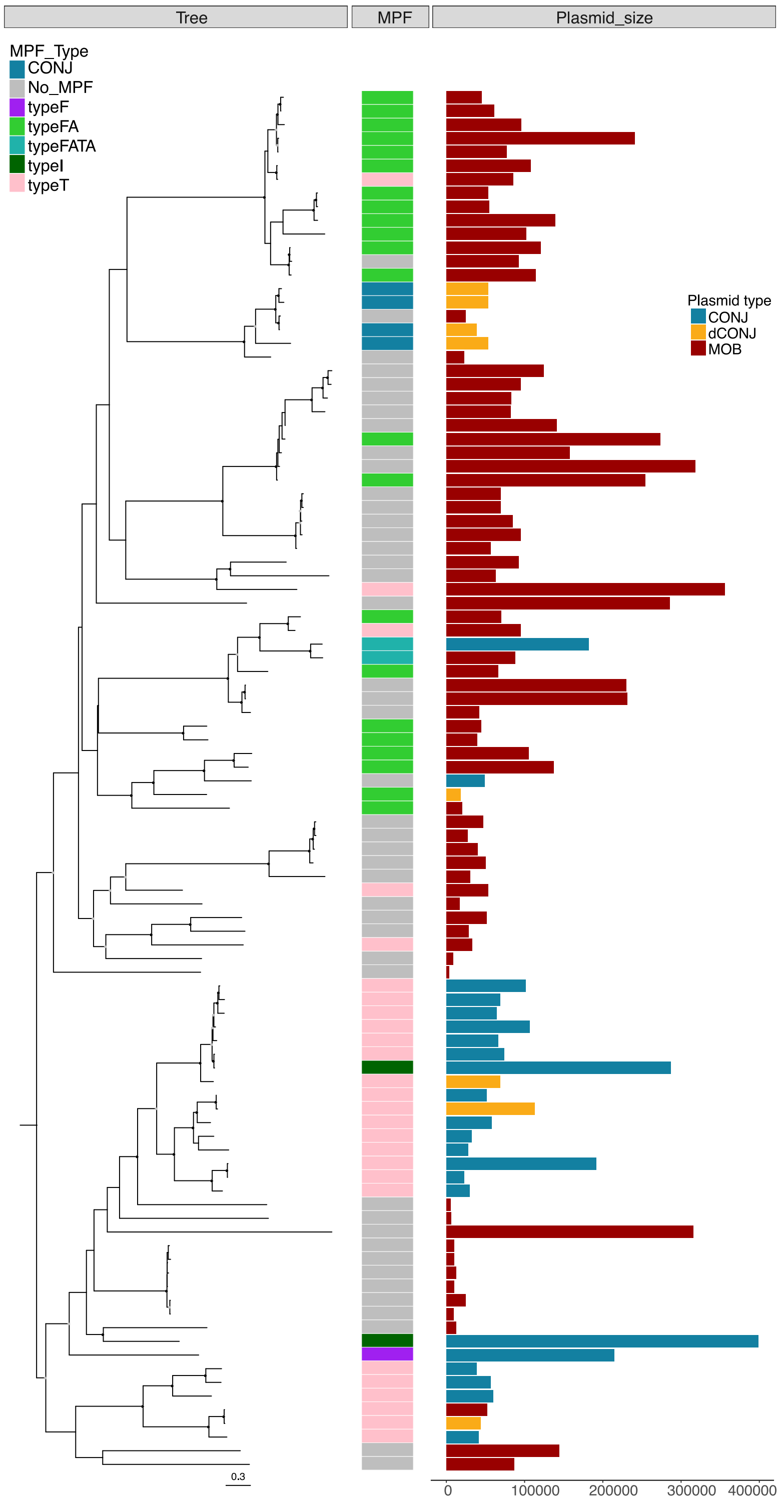


**Figure S18: Phylogenetic tree of MOB$_C$ relaxases.** Ultrafast bootstrap values superior to 75 are shown with a light grey circle and values superior to 90 with a black circle. The trees were rooted using the mid-point root. The phylogenetic tree was built using 101 MOBC proteins with maximum likelihood with IQTree (model LG+R5 and 1000 ultrafast bootstraps).

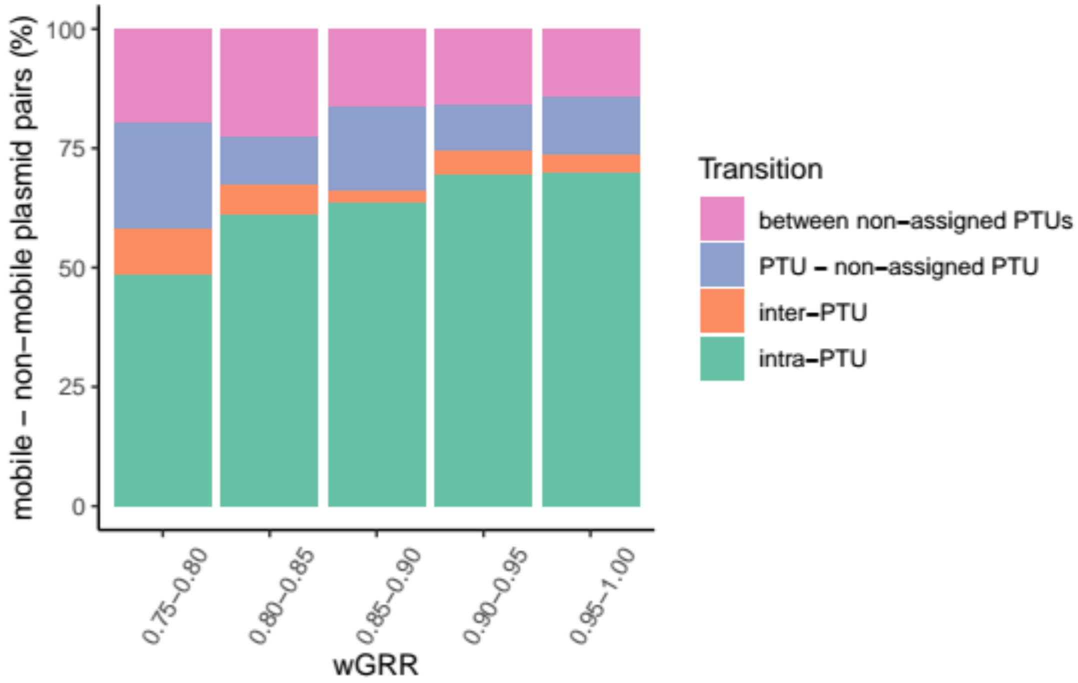


**Figure S19: PTU transitions between mobile and non-mobile plasmid pairs.** Pairs of mobile-non mobile plasmids with wGRR > 0.75 were analyzed regarding their PTUs. In green, mobile - non-mobile pairs whose members belong to the same PTU. In orange, mobile - non-mobile pairs whose members belong to different PTUs. In blue, mobile - non-mobile pairs in which a member belongs to an assigned PTU and the other one to a non-assigned PTU. In pink, mobile - non-mobile pairs whose members belong to non-assigned PTUs.
